# Monophyly of Diverse Bigyromonadea and their Impact on Phylogenomic Relationships Within Stramenopiles

**DOI:** 10.1101/2021.11.17.469027

**Authors:** Anna Cho, Denis V. Tikhonenkov, Elisabeth Hehenberger, Anna Karnkowska, Alexander P. Mylnikov, Patrick J. Keeling

## Abstract

Stramenopiles are a diverse but relatively well-studied eukaryotic supergroup with considerable genomic information available (Sibbald and Archibald, 2017). Nevertheless, the relationships between major stramenopile subgroups remain unresolved, in part due to a lack of data from small nanoflagellates that make up a lot of the genetic diversity of the group. This is most obvious in Bigyromonadea, which is one of four major stramenopile subgroups but represented by a single transcriptome. To examine the diversity of Bigyromonadea and how the lack of data affects the tree, we generated transcriptomes from seven novel bigyromonada species described in this study: *Develocauda condao*, *Develocanicus komovi*, *Develocanicus vyazemskyi*, *Cubaremonas variflagellatum*, *Pirsonia chemainus*, *Feodosia pseudopoda*, and *Koktebelia satura*. Both maximum likelihood and Bayesian phylogenomic trees based on a 247 gene-matrix recovered a monophyletic Bigyromonadea that includes two diverse subgroups, Developea and Pirsoniales, that were not previously related based on single gene trees. Maximum likelihood analyses show Bigyromonadea related to oomycetes, whereas Bayesian analyses and topology testing were inconclusive. We observed similarities between the novel bigyromonad species and motile zoospores of oomycetes in morphology and the ability to self-aggregate. Rare formation of pseudopods and fused cells were also observed, traits that are also found in members of labyrinthulomycetes, another osmotrophic stramenopiles. Furthermore, we report the first case of eukaryovory in the flagellated stages of Pirsoniales. These analyses reveal new diversity of Bigyromonadea, and altogether suggest their monophyly with oomycetes, collectively known as Pseudofungi, is the most likely topology of the stramenopile tree.

## 1. Introduction

Stramenopiles (= Heterokonts) are one of the well-characterized members of the eukaryotic supergroup SAR (**S**tramenopila, **A**lveolate, **R**hizaria) (Keeling and Burki, 2019). Stramenopiles are very diverse, comprising photoautotrophs (i.e. heterokont algae in ochrophytes), osmotrophic oomycetes and labyrinthulomycetes with a motile zoospore life-stage (e.g. *Phytophthora* sp*., Pythium sp.,* and labyrinthulids), and, free-living phagotrophic opalozoans (e.g. *Cafeteria roenbergensis, Cantina marsupialis*) that occupy a broad range of environments (Cavalier-Smith and Chao, 2006; Cavalier-Smith and Scoble, 2013a; Kolodziej and Stoeck, 2007; Stiller et al., 2009; Tsui et al., 2009). Stramenopiles can be largely classified into two major groups: Gyrista consisting of Ochrophyta, Oomycota, and Bigyromonadea; and Bigyra consisting of Sagenista and Opalozoa. A single species *Platysulcus tardus,* has also recently been shown to be a basal stramenopile (Thakur et al., 2019). While there is a lot of genomic data from stramenopiles, only a handful comes from phagoheterotrophs (Mitra et al., 2016), despite them representing much of the diversity as well as being key outstanding problems in resolving controversies in stramenopiles phylogeny (Burki et al., 2016; Derelle et al., 2016; Leonard et al., 2018; Shiratori et al., 2017, 2015).

One such clade is the subphylum Bigyromonadea, which was proposed to include the class Developea (Aleoshin et al., 2016) and order Pirsoniales (Cavalier-Smith, 1998). The monophyly of the Bigyromonadea is essentially untested, since only small subunit rRNA (SSU) data are known from all but a single species (the exception being *Developayella elegans*, for which a transcriptome is available), and the two groups never branch together in SSU phylogenies (Aleoshin et al., 2016; Cavalier-Smith and Chao, 2006; Kühn et al., 2004; Weiler et al., 2020).

Developea are marine bacterivores, including *Developayella elegans* (Leipe et al., 1996; Tong, 1995) and *Mediocremonas mediterraneus* (Weiler et al., 2020), and the marine eukaryovore *Develorapax marinus* (Aleoshin et al., 2016). Pirsoniales are parasites of other microbes, including *Pirsonia guinardiae* (Schnepf et al., 1990) and *P. punctigera* (Schweikert and Schnepf, 1997). These parasites deploy a pseudopod to squeeze through the frustule girdles of their diatom host, while the main cell body (auxosome) stays outside of the host. The invading pseudopod then phagocytoses the host cytoplasm or chloroplasts forming a trophosome (food vacuole), which is then transported out to the auxosome (Kühn et al., 2004; Schnepf et al., 1990).

The relationship of both groups to other stramenopiles is uncertain, and both have led to hypotheses about the evolution of other related groups. For example, the eukaryovory of *D. marinus* and its placement in rRNA trees has led to the hypothesis that it represents a model for a phagoheterotrophic ochrophyte ancestor (Aleoshin et al., 2016), however its position in the tree varies between grouping with ochrophytes (Leonard et al., 2018) or oomycetes (Noguchi et al., 2016; Thakur et al., 2019). Pirsoniales have also been found branching as sister to ochrophytes based on SSU rRNA trees (Aleoshin et al., 2016; Kühn et al., 2004), although once again not consistently and without strong support.

To test for the monopoly of bigyromonads and more thoroughly examine their relationship to other stramenopiles, we significantly increased the diversity of genomic data from the group by adding transcriptomes from seven newly discovered species belonging to Pirsoniales (*Pirsonia chemainus* nom. prov., *Koktebelia satura* nom. prov., and *Feodosia pseudopoda* nom. prov.) and Developea (*Develocanicus komovi* n. gen. n. sp., *Develocanicus vyazemskyi* n. sp., *Develocauda condao* n. gen. n. sp., and *Сubaremonas variflagellatum* n. gen. n. sp.). The inferred 247-gene phylogenomic tree, reconstructed with various methods, recovered for the first time the monophyly of the Bigyromonadea. Maximum likelihood (ML) recovered a robust monophyly of Bigyromonadea and oomycetes, while Bayesian inference and topology testing were inconclusive. We describe several new features of the seven bigyromonads, and noted their resemblance with oomycete zoospores, and report the first observation of eukaryovory in the flagellated stages of Pirsoniales. Overall, these findings indicate bigyromonada and ooymcetes are most likely sisters, and suggest potential ancestral state of the oomycetes resembling bigyromonada, including their ability to form auto-aggregates (=self-aggregates) (Galiana et al., 2008; Hickman, 1970; Ko and Chase, 1973) and phagoheterotrophy.

## 2. Materials and Methods

### 2.1 Sample collection, identification, and library preparation

Strain Сolp-23 (*Develocanicus komovi*) was obtained from the black volcanic sand on the littoral zone of Maria Jimenez Beach (Playa Maria Jiménez), Puerto de la Cruz, Tenerife, Spain, October 20, 2014. Strains Colp-30 (*Develocanicus vyazemskyi*) and Chromo-1 (*Koktebelia satura*) were isolated from the near shore sediments on the littoral zone near T.I. Vyazemsky Karadag Scientific Station, Crimea, May 2015. Strain Chromo-2 (*Feodosia pseudopoda*) was obtained from the near shore sand on the littoral zone of the beach in the settlement Beregovoye, Feodosiya, Crimea, June 24, 2017. Strain Colp-29c (*Develocauda condao*) was isolated from the near shore sediments on the north-east part of Con Dao Island, South Vietnam, May 4, 2015. Strains ‘*Pirsonia*-like’ (*Pirsonia chemainus*) and Dev-1 (*Сubaremonas variflagellatum*) were obtained from a sea water samples taken in the Strait of Georgia, British Columbia, Canada (123° 28’50’’ W, 49°10’366’’ N) at 70 m and 220 m depths, respectively using a Niskin bottle, June 13, 2017.

The samples were examined on the third, sixth and ninth day of incubation in accordance with methods described previously (Tikhonenkov et al., 2008). *Procryptobia sorokini* strain B-69 (IBIW RAS), feeding on *Pseudomonas fluorescens*, was cultivated in Schmaltz-Pratt’s medium at a final salinity of 20‰, and used as a prey for clones Colp-23, Colp-29c, Colp-30, Chromo-1, Chromo-2, and ‘*Pirsonia*-like’ (Tikhonenkov et al., 2014). Bacterivorous strain Dev-1 was propagated on the *Pseudomonas fluorescens*, which was grown in Schmaltz-Pratt’s medium. Strains Colp-23, Colp-29c, and Dev-1 are currently being stored in a collection of live protozoan cultures at the Institute for Biology of Inland Waters, Russian Academy of Sciences, however, strains Colp-30, Chromo-1, Chromo-2, and ‘*Pirsonia*-like’ perished after several months to one year of cultivation.

Studied isolates were identified using a combination of microscopic and molecular approaches. Light microscopy observations were made using a Zeiss AxioScope A.1 equipped with a DIC water immersion objective (63x) and an AVT HORN MC-1009/S analog video camera. The SSU rRNA genes (GenBank accession numbers: XXXXXX, XXXX…) were amplified by polymerase chain reaction (PCR) using the general eukaryotic primers EukA-EukB (for strains Colp-23, Colp-30, ‘*Pirsonia*-like’), PF1-FAD4 (Chromo-1), 18SFU-18SRU (Chromo-2, Dev-1), 25F-1801R (Colp-29c) (Cavalier-Smith et al., 2009; Keeling, 2002; Medlin et al., 1988; Tikhonenkov et al., 2016). PCR products were subsequently cloned (Colp-23, Colp-30, Chromo- 2, ‘*Pirsonia*-like’) or sequenced directly (Chromo-1, Dev-1, Colp-29c) using Sanger dideoxy sequencing.

For cDNA preparation, cells grown in clonal laboratory cultures were harvested when the cells had reached peak abundance (strains Colp-23, Col-30, Colp29c, Chromo-1, Dev-1) and after the majority of the prey had been eaten (for eukaryovorous strains Colp-23, Col-30, Colp29c, Chromo-1). Cells were collected by centrifugation (1000 x *g*, room temperature) onto the 0.8 µm membrane of a Vivaclear mini column (Sartorium Stedim Biotech Gmng, Cat. No. VK01P042). Total RNA was then extracted using a RNAqueous-Micro Kit (Invitrogen, Cat. No. AM1931) and reverse transcribed into cDNA using the Smart-Seq2 protocol (Picelli et al., 2014), which uses poly-A selection to enrich mRNA. Additionally, cDNA of Colp-29c was obtained from 20 single cells using the Smart-Seq2 protocol (cells was manually picked from the culture using a glass micropipette and transferred to a 0.2 mL thin-walled PCR tube containing 2 µL of cell lysis buffer – 0.2% Triton X-100 and RNase inhibitor (Invitrogen)). The same ‘single cell’ transcriptomic approach was applied for strains Chromo-2 and ‘*Pirsonia*-like’, which never consumed the prey completely. Sequencing libraries were prepared using NexteraXT protocol and sequencing was performed on an Illumina MiSeq using 300 bp paired-end reads. Additionally, Chromo-1 transcriptome sequencing was performed on the Illumina HiSeq platform (UCLA Clinical Microarray Core) with read lengths of 100 bp using the KAPA stranded RNA-seq kit (Roche) to construct paired-end libraries. Raw reads are available in the NCBI Short Read Archive (SRA) (Bioproject number: XXXXXX, SRAXXX-XXX).

### 2.2 Small-subunit phylogenetic tree reconstruction

SSU rRNA sequences were identified from the seven new assembled transcriptomes using Barrnap v0.9 (Seemann, 2007) and compared with the SSU sequences obtained with Sanger sequencing, and the longer sequences were used for further analysis.

After an initial BLASTn search of the SSU rRNA sequences against the non-redundant NCBI database to confirm stramenopile identities, the SSU sequences were aligned using MAFFT v7.222 (Katoh and Standley, 2013) with previously compiled SSU datasets (Aleoshin et al., 2016; Yubuki et al., 2015). Additionally, SSU sequences of the other stramenopile taxa that were included in the multi-gene phylogenomic dataset and other closely related taxa were included in Fig. 3. Furthermore, to show the diversity of uncultured Gyrista and provide possible directions for future sampling efforts, environmental sequences of stramenopiles that are closely related to Pirsoniales and Developea were added in Fig. S4. After trimming using trimAl v.1.2rev59 (-gt 0.3, -st 0.001) (Capella-Gutiérrez et al., 2009), the SSU phylogenetic trees were reconstructed based on 1650 sites and 92 taxa for Fig. 3, and 1665 sites and 107 taxa for Fig. 4S, using IQ-TREE v1.6.12 (Nguyen et al., 2015) 1000 ultrafast bootstrap (UFB) under Bayesian information criterion (BIC): TIM2+R6 selected by ModelFinder (Kalyaanamoorthy et al., 2017) implemented in IQ-TREE.

### 2.3 Transcriptome processing, assembly, and decontamination

Raw sequencing reads were assessed for quality using FastQC v0.11.5 (Andrews, 2010) and remnant transposase-inserts from the library preparation were removed. The reads were assembled using Trinity-v2.4.0 with *–trimmomatic* option to remove NexteraXT adaptors, Smart-Seq2 IS-primer, and low quality leading and trailing ends (quality threshold cut-off:5) (Bolger et al., 2014; Grabherr et al., 2011). To identify contaminants, assembled reads were searched against the NCBI nucleotide database using megaBLAST (Basic Local Alignment Search Tool) (Altschul et al., 1990), followed by diamond BLASTX against a UniProt reference proteome (Bateman et al., 2021). To visualize the contig sizes, coverage, and remove bacterial, archaeal, and metazoan contaminants, BlobTools v1.0 (Laetsch and Blaxter, 2017) was used.

PhyloFlash v3.3b2 (Gruber-Vodicka et al., 2020) was used in parallel to confirm identified contaminants and coverage based on SILVA v138 SSU database (Quast et al., 2013). To remove sequences from the prey, *Procryptobia sorokini,* which was used in the cultures of *Pirsonia chemainus*, *Koktebelia satura*, *Feodosia pseudopoda*, *Develocanicus komovi*, *D. vyazemskyi*, and *Develocauda condao*, the assembled reads were searched against the *P. sorokini* transcriptome using BLASTn in which the contigs with ≥95% sequence identity were removed from the assembled reads. To predict open reading frames (ORFs) and coding genes, TransDecoder v5.5.0 (Haas, 2015) was used and the longest ORFs were annotated using BLASTP search against UniProt database. To estimate the completeness of each of the assembly, BUSCO v4.0.5 (Simão et al., 2015) with eukaryotic database was used.

### 2.4 Phylogenomic matrix construction and ortholog identification

To better represent each stramenopile (sub)group in the phylogenomic reconstruction, recently published and publicly available (Broad Institute and Japan Agency for Marine-Earth Science and Technology; JAMSTEC) additional 27 stramenopile genomic or transcriptome data (de Vargas et al., 2015; Hackl et al., 2020; Keeling et al., 2014; Leonard et al., 2018; Noguchi et al., 2016; Seeleuthner et al., 2018; Thakur et al., 2019; Wawrzyniak et al., 2015) were obtained and analyzed along with the seven new transcriptomes (Table S1). The updated stramenopile dataset including all the newly added transcriptomes in this study were compiled to the existing gene-set described below. Using BLASTP, the predicted coding genes from each transcriptome were searched against 263 gene-sets (orthologs), each consisting of compiled genes from major supergroups of protists, fungi, and holozoans (Burki et al., 2016; Hehenberger et al., 2017). The search results were filtered with an e-value threshold of 1e-20 with >50% query coverage, followed by read trimming based on UniProt search results. Each gene-set was aligned using MAFFT-L-INS-i v.7222 and trimmed using trimAL v1.2rev59 (-gt 0.8). To identify orthologs from the newly added transcriptomes aligned to the corresponding 263 gene-sets, 263 gene-trees were built using Maximum-likelihood (ML) estimation with IQ-TREE v1.6.12 under the LG+I+G4 model and 1000 ultrafast bootstrap (UFB) approximation. Each gene tree was manually screened in FigTree v1.4.4 and searched against BLAST nr-database for paralogs and contaminants (e.g., long branching sequences or sequences nested within bacterial, opisthokont and other distantly related clades), which were subsequently removed from each of the gene-set alignment. To increase ortholog coverage from the added transcriptomes, fragmented orthologs were manually combined only when the fragments were positioned in the same clade in a given gene tree, covered different regions of a gene, and if there was an overlapping region present, permitting up to two mismatches. The 263 gene-sets containing the selected orthologs of the newly added transcriptomes and 27 newly published stramenopile data were aligned using two approaches and compared by reconstructing two phylogenomic trees. In the first approach (**approach 1**), the sequences were aligned by using MAFFT L-INS-i v.7.222 and trimmed via trimAL v1.2rev59 (-gt 0.8). In the second approach (**approach 2**), the sequences were filtered using PREQUAL (Whelan et al., 2018) to remove non-homologous regions generated due to poor transcriptome quality or assembly errors. The filtered sequences were then aligned using MAFFT G-INS-i (--allowshift and --unalignlevel 0.6 option) and processed for further filtering using Divvier (-mincol 4 and -divvygap option) (Ali et al., 2019) to identify statistically robust pairwise homology characters. The filtered gene-sets were then soft-trimmed using trimAL (-gt 0.1). The two dataset generated by two different filtering and alignment methods were separately processed using SCaFoS v1.2.5 (Roure et al., 2007), by removing gene-sets that have ≥40% missing amino acid positions in the alignment. The resulting 247 gene-set was concatenated into a phylogenomic matrix comprising 75,798 amino acid (aa) sites from 76 taxa for approach 1. For the PREQUAL/Divvier processed data (approach 2), the same 247 gene-sets were concatenated in a phylogenomic matrix comprising 101,314 aa sites from the same 76 taxa.

**Table 1.** Approximately unbiased (AU) tests on tree constraints based on approach 1 dataset.

| Approach 1 (MAFFT L-INS-i and trimAl -g 0.8) |  |  |  |
| --- | --- | --- | --- |
| Constrained Tree | p-AU | logL | ΔlogL |
| Unconstrained ML tree | 0.78 | -3935691.338 | 0 |
| ML tree | 0.759 | -3935691.338 | 0.00089261 |
| Chain 1 (C+S+Pi),(R+P+X+E) | 0.0541 | -3935828.083 | 136.75 |
| Chain 1 Modified (Bigyromonada+oomycetes) | 0.267 | -3935763.845 | 72.508 |
| Chain 2 (C+S+Pi+E),(R+P+X) | **0.0297** | -3935859.39 | 168.05 |
| Chain 2 Modified (Bigyromonada+oomycetes) | 0.0924 | -7871604.549 | 108.01 |
| Chain 3 (R+P+X+E),(C+S) | **0.0119** | -3935874.998 | 183.66 |
| Chain 3 Modified (Bigyromonada+oomycetes) | 0.0717 | -3935805.205 | 113.87 |
| Chain 4 (C+S+E),(R+P+X) | **0.0186** | -3935860.003 | 168.67 |
| Chain 4 Modified (Bigyromonada+oomycetes) | 0.108 | -3935765.741 | 74.404 |

### 2.5 Phylogenomic tree reconstruction, fast-evolving site removal, and topology test

The ML tree for the concatenated phylogenomic matrix was inferred using IQ-TREE v.1.6.12 under the empirical profile mixture model, LG+C60+F+G4 (Quang et al., 2008). The best tree under this model was used as a guide tree to estimate the “posterior mean site frequencies” (PMSF). The PMSF method allows the conduction of non-parametric bootstrap analyses under complex models on large data matrices and was shown to mitigate long-branch attraction artifacts (Wang et al., 2018). This LG+C60+F+G-PMSF model was then used to re-estimate the ML tree and for a non-parametric bootstrap analysis with 100 replicates. For Bayesian inference, CAT-GTR mixture model with four gamma rate categories was used with PhyloBayes-MPI v.20180420 (Lartillot et al., 2009; Lartillot and Philippe, 2004), only for the dataset processed with approach 1. To estimate posterior probability, four independent Markov Chain Monte Carlo (MCMC) chains were run simultaneously for minimum 10,000 cycles. After discarding the first 2000 burn-in points, consensus posterior probability and topology were computed by subsampling every second tree. Convergence of the four chains were tested by calculating differences in bipartition frequencies (bpcomp) with a threshold maxdiff however, no chains converged (maxdiff=1).

Site-specific substitution rates were inferred using the -wsr option as implemented in IQ-TREE, under the LG+C60+F+G4 substitution model. Increments of the top 5% fastest evolving sites were removed (Irwin, 2021) from the phylogenomic matrix until exhaustion, defined as the point when the bootstrap support value significantly began to drop and the topology became unstable (50%; 37,899 sites). Each incremental phylogenomic matrix was analyzed using IQ-TREE for ML estimation using LG+C60+F+G4 and 1000 UFB. All fast-evolving species removal and sites tests were conducted on the dataset processed with approach 1.

Approximately unbiased (AU) tests (Nguyen et al., 2015; Shimodaira, 2002) were performed on set of phylogenomic trees constructed based on the 247 gene-sets generated by the first approach (i.e., MAFFT L-INS-i and trimAL with -gt 0.8) and the second approach (i.e., PREQUAL/Divvier), separately. The set of trees includes the two ML trees generated under LG+C60+F+G4(+PMSF) with 1000UFB (100STB), four consensus trees of MCMC chains, and other hypothetical constrained trees as listed as “Chain modified” in Table 1.

## 3. Results

### 3.1 Multi-gene phylogenomic analysis

The concatenated phylogenomic matrix was composed of 68 stramenopiles and eight alveolates (outgroup) with 247 aligned genes totaling 75,798 positions for approach 1, and 101,314 positions for approach 2. The average missing sites and genes were 22% and 19%, respectively (Fig. 1). The amount of missing data varied among the seven new transcriptomes. Chromo-1 had nearly complete data (5% missing sites and 6% missing genes) while Colp-29c had 21% missing sites and 12% missing genes. Colp-23 and Chromo-2 had the highest amount of missing data (75% missing sites and 57% genes for Chromo-2 and, 83% and 76% for Colp- 23). The ML phylogenomic tree generated under LG+C60+F+G4+PMSF with STB estimation from the two approaches is shown in Figure 1, with the tree topology representing the dataset generated from approach 1 (i.e., MAFFT L-INS-i and trimAL with -gt 0.8). The tree topology representing the dataset generated from approach 2 (i.e., Prequal/Divvier) is shown in Figure S1. The tree topology is almost identical between the two, except the position of sub-clades in ochrophytes; for example, the positions of Chrysophyceae + Synurophyceae and Raphidophyceae + Phaeophyceae + Xanthophyceae + Eustigmatophaceae are swapped in the two trees (Fig. 1; Fig. S1).

**Figure 1.**
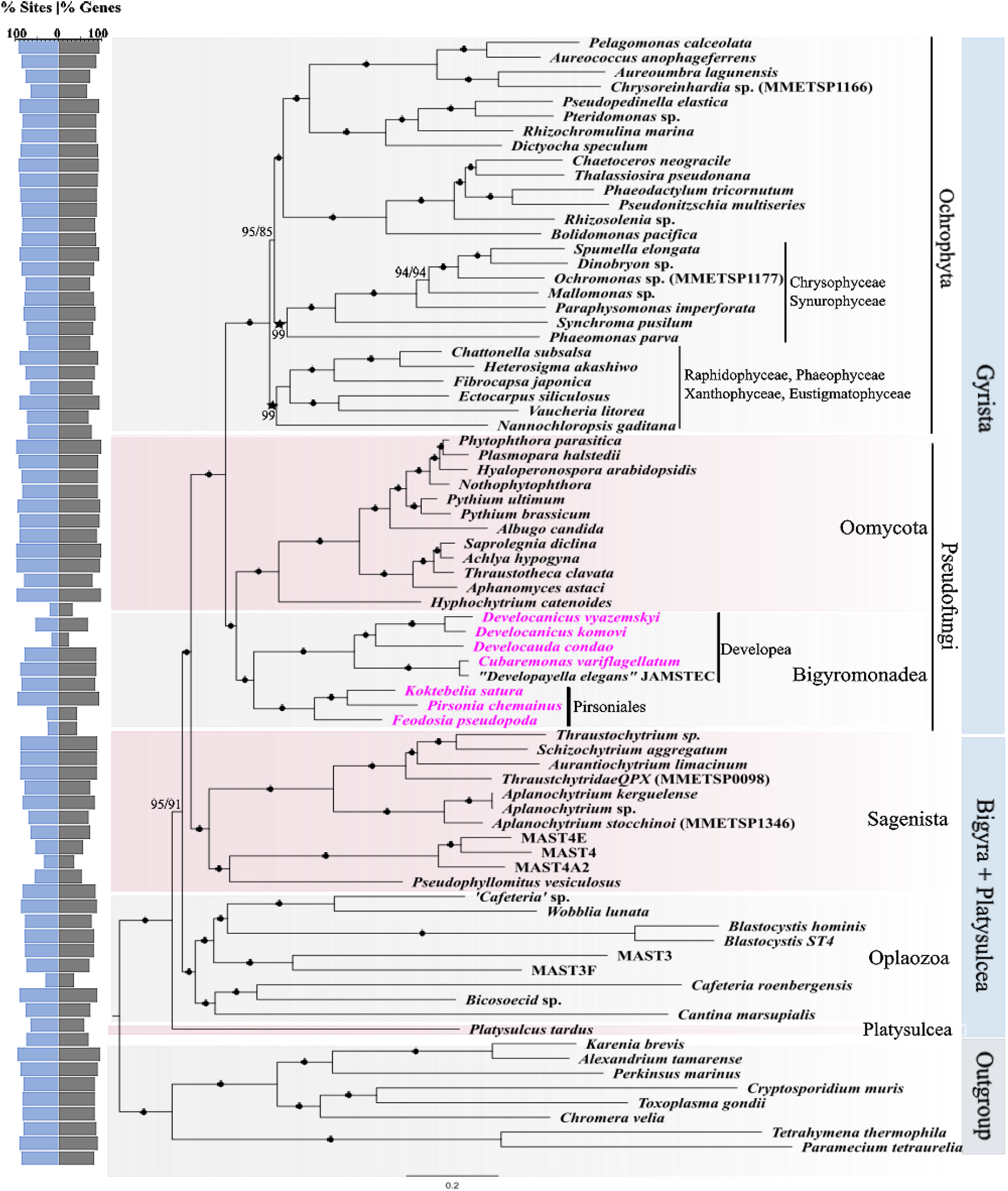
Multi-gene phylogenomic tree of stramenopiles with the seven new transcriptomes (pink) added to Gyrista, consisting of the concatenated alignments of 247 genes of 76 taxa. The tree was reconstructed using the Maximum-likelihood (ML) analysis, under the site-heterogenous model, LG+C60+F+G4+PMSF, implemented in IQ-Tree. Branch support was calculated using non-parametric PMSF 100 standard bootstrap (STB). Branches with ≥99% STB for both approaches are marked with black bullets while others are labelled as “Approach 1 STB/Approach 2 STB”. The topology of the trees generated from the two approaches were the same except the positions of Raphidophyceae, Phaeophyceae, Xanthophyceae + Eustigmatophacea and Chrysophyceae+Synurophceae, which were swapped in the tree reconstructed based on the dataset processed using approach 2 (i.e., Prequal/Divvier method); denoted by star symbols (Fig. S1). The percent sites (blue) and genes (grey) present for each transcriptome is depicted on the back-to-back bar plot on the left.

The newly added transcriptomes of the seven new species formed the robust monophyletic bigyromonada with either dataset (approach 1 and approach 2; Fig. 1; Fig. S1): *Develocanicus komovi*, *D*. *vyazemskyi*, *Develocauda condao,* and *Сubaremonas variflagellatum* forming a Developea clade (100% STB), while Pirsoniales is composed of *Pirsonia chemainus*, *Koktebelia satura*, and *Feodosia pseudopoda* (100% STB). The ML tree also recovered monophyly of the bigyromonada and oomycetes with 100% STB support (Fig. 1). The monophyly of Gyrista was strongly supported, with Sagenista (Labyrinthulomycetes and Eogyrea) forming a sister clade to it, resulting in a paraphyletic Bigyra. Platysulcea formed a sister clade to rest of the stramenopiles with a moderate support (91%/95% STB) (Fig.1; Fig. S1).

Bayesian analyses recovered a conflicting topology for the bigyromonada, which formed a sister-clade to ochrophytes in all four consensus trees generated (Fig. S2). Additionally, the topology within ochrophytes was inconsistent, preventing convergence. However, the monophyly of bigyromonada + ochrophytes was rejected by approximately unbiased (AU) tests in three of the four consensus trees. AU test failed to reject the chain 1 consensus tree at a confidence interval of 95% (p-AU ≥ 0.05) (Fig. S2). Interestingly, the sub-clade topology of ochrophytes in chain 1 is the same as in the ML phylogenomic tree generated using the approach 1 (Fig. 1; Fig. S2). When the AU tests were repeated on hypothetically constrained trees where bigyromonada + oomycetes were monophyletic but the rest of the topology was unchanged for each of the MCMC chains, the tests failed to reject the monophyly of bigyromonada + oomycetes (Table 1). Rejection of bigyromonada + ochrophytes was also observed in constrained trees when the AU test was repeated on the dataset processed with approach 2 (Table S2). To evaluate the effect of fast-evolving sites, bootstrap support and topology were compared among the ML trees that were reconstructed with increments of 5% fast-evolving sites removed from the dataset processed with approach 1. The topologies of the phylogenomic tree were maintained while the UFB support for Platysulcea increased up to 97% (Fig. 2). To account for possible artefacts due to long-branching attraction of fast-evolving species, tree reconstruction was repeated after removing *Cafeteria roenbergensis*, the two *Blastocystis* species, and *Cantina marsupialis*. The monophyly of bigyromonada and oomycetes was recovered with 85% UFB, however the topology of Bigyra became unresolved with weak support for its monophyly (Fig. S3)

**Figure 2.**
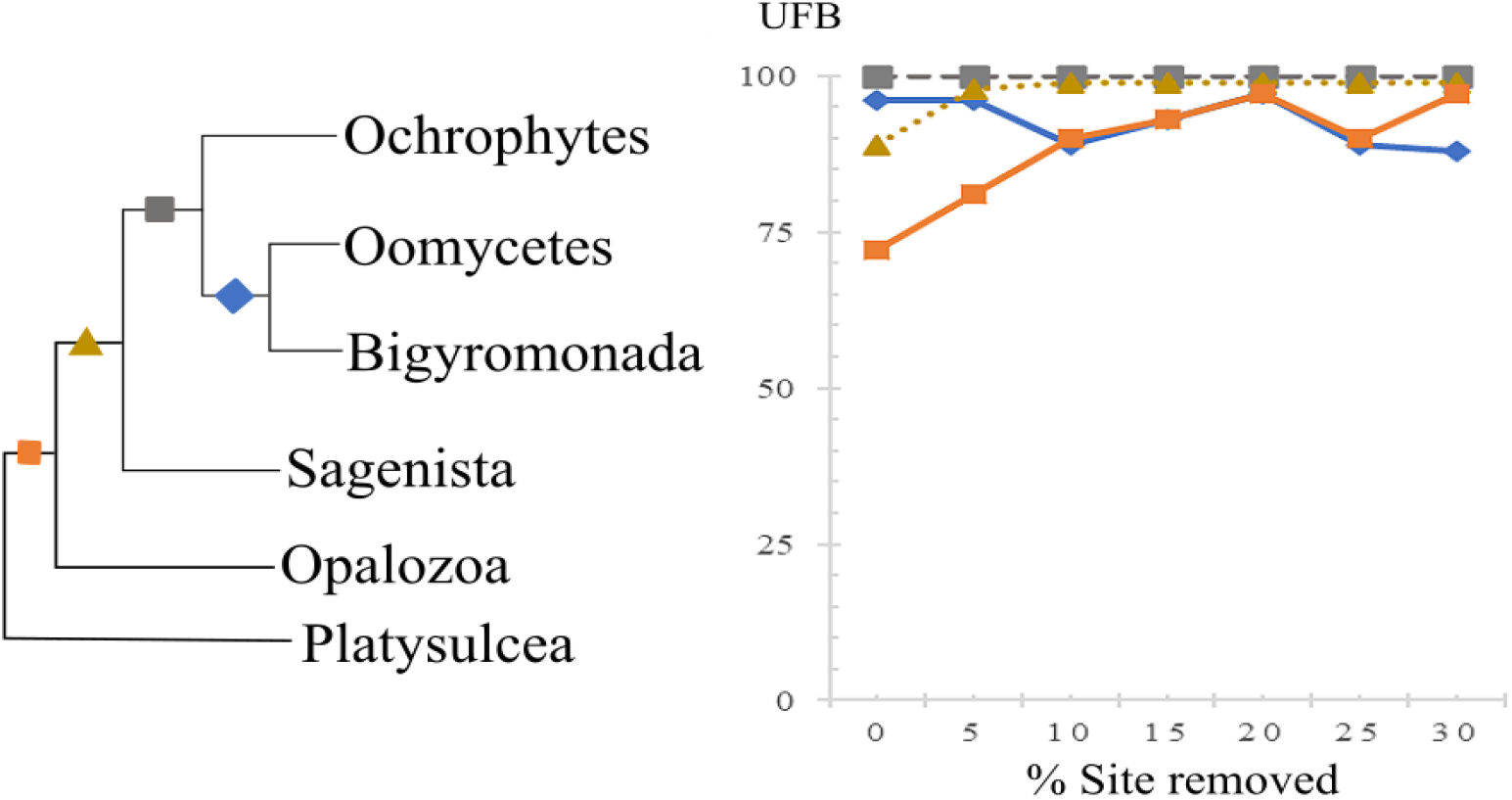
Summary of ultrafast bootstrap (UFB) with incremental removal of fast-evolving sites, based on the dataset processed with approach 1. Schematic representation the stramenopiles ML tree (left) with each branch marked with different shapes and colours. The line plot (right) showing the change in UFB for each branch when fast-evolving sites were incrementally removed by 5%. The monophyly of Gyrista shows full support throughout while the UFB increases incrementally for ‘Sagenista’ and ‘Platysulcea’.

### 3.2 Small-subunit ribosomal RNA gene tree reveals two different species assigned as *Developayella*

As shown previously, the SSU rRNA phylogenetic tree recovered the bigyromonada as paraphyletic group, with the Pirsoniales (*Pirsonia chemainus*, *Koktebelia satura*, and *Feodosia pseudopoda*) forming a sister clade to ochrophytes (92% UFB) while the Developea clade was recovered as sister to oomycetes (Fig. 3). Within the Developea clade, in addition to the SSU rRNA sequences obtained from *Сubaremonas variflagellatum* and the JAMSTEC *Developayella elegans* transcriptome, we included three publicly available SSU rRNA sequences assigned as *Developayella* spp.: Accession ID U37107 (Leipe et al., 1996; Tong, 1995), MT355111.1 (Unpublished) and JX272636.1 (Del Campo et al., 2013): (note: although JX272636.1 is assigned as “Cf. *Developayella* sp.” in GenBank, it was recently re-assigned as *Mediocremonas mediterraneus* (Weiler et al., 2020)). Interestingly, the SSU rRNA sequences of the four “*Developayella*” fell into two separate groups, indicating two different species (and genera) were assigned as *Developayella elegans*; sub-clade I consisted of *Developayella elegans* U37107, *Developayella* sp. MT355111.1, *Develocanicus komovi*, *D*. *vyazemskyi*, and *Develocauda condao,* while sub-clade II consisted of *M. mediterraneus* (JX272636.1 and MT918788.1), JAMSTEC *Developayella elegans*, and *Сubaremonas variflagellatum* (Fig. 3). The SSU rRNA sequence similarity between the two sub-clade I *Developayella* species (U37107 and MT355111.1) is 98.987%, between the two species (JAMSTEC *D. elegans* and *Сubaremonas variflagellatum*) in sub-clade II 97.528% and between the originally described *Developayella elegans* U37107 and *Сubaremonas variflagellatum* 91.143%.

**Figure 3.**
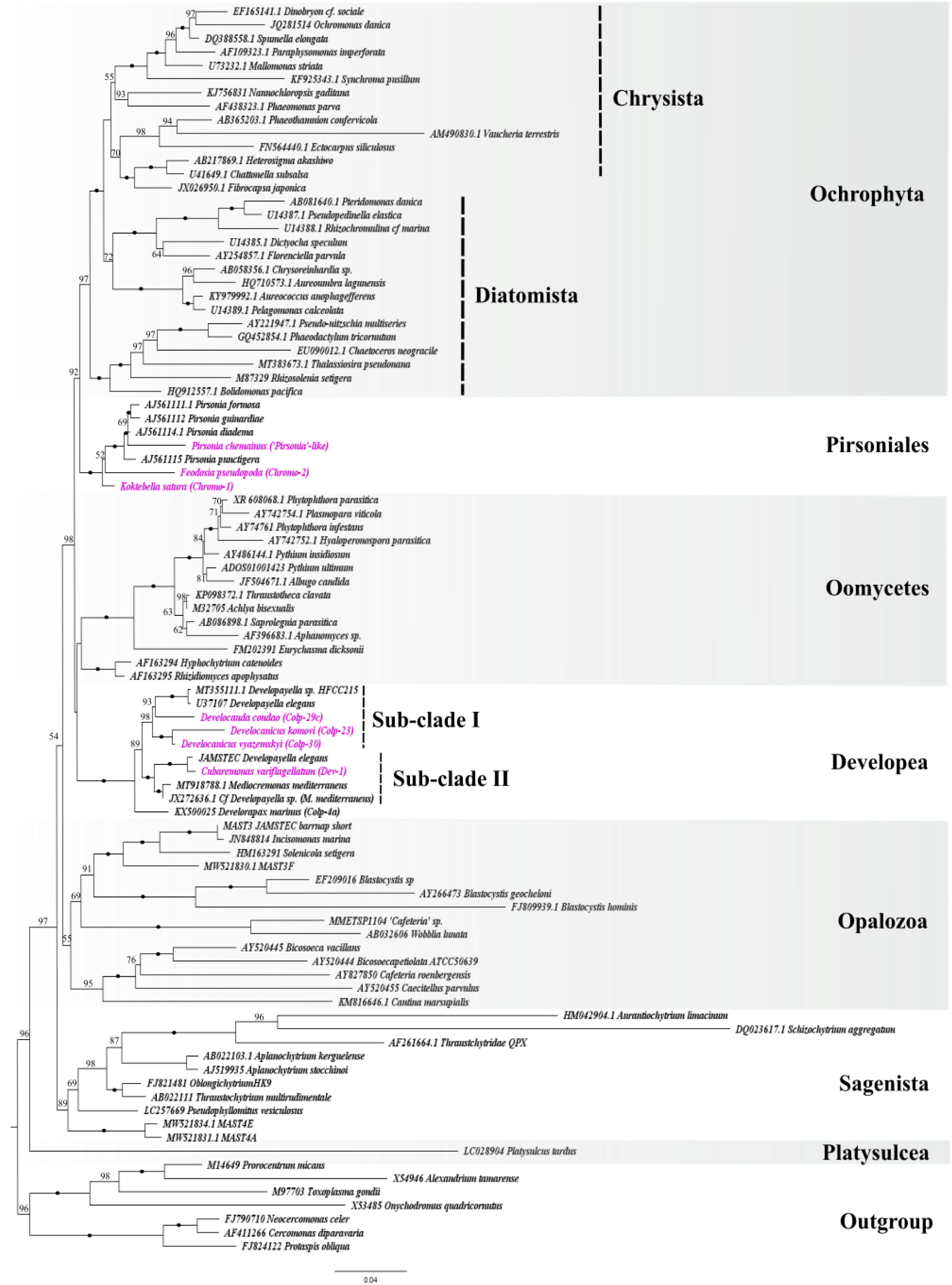
ML tree reconstructed from a 18S rRNA gene alignment of 92 taxa (1650 sites), under BIC: TIM2+R6 with 1000 UFB. Branch support with ≥99% UFB is marked with black bullets while the values less than 50% are not shown. The seven new species described in this study are marked as pink: Pirsoniales forming a sisterhood with Ochrophytes and Developea forming a sister clade to Oomycetes. Within Developea, two previously assigned *Developayella* species (JAMSTEC transcriptome and the U37107 SSU rRNA sequence) are split into two sub-clades, in which the four novel Developea species are positioned.

### 3.3 Morphology of the novel species

Developea Karpov et Aleoshin 2016

#### Develocanicus vyazemskyi (Fig. 4A, B) and Develocanicus komovi (Fig. 4C–M)

Free-swimming naked eukaryovorous heterokont flagellates. The shape of the cell is irregularly flattened ellipse, where the dorsal side is more convex, and the ventral side is flatter. Two species differ in size, *Develocanicus vyazemskyi* (Colp-30) is larger and rounder, 7.4 - 12.5 μm long, 4.8 - 9.2 μm wide, typical dimension ranging 9.2 х 7.0 μm. *Develocanicus komovi* (Colp-23) is slightly smaller, with the length 5.4 - 10 μm, width 3.8 - 7.4 μm and a typical dimension of 7.1 х 5.1 μm.

Cell possesses two non-acronematic heterodynamic flagella of unequal lengths (Fig. 4A-D, F, I, J). The posterior flagellum is two times longer than the cell, the anterior flagellum is approximately 1 - 1.5 times longer. Flagella emerge from a prominent ventral depression (Fig. 4A-D) which passes into a shallow wide groove (Fig. 4E) along the entire length of the cell. Cells predominantly exhibit active and quick swimming without rotation. During swimming, the posterior flagellum is directed backward and straight, running along the ventral depression of the cell. The anterior flagellum beats rapidly and is directed forward while slightly curved. In non-motile cells, both flagella are directed backward, beating in a slow sinusoidal wave (Fig. 4G, J).

**Figure 4.**
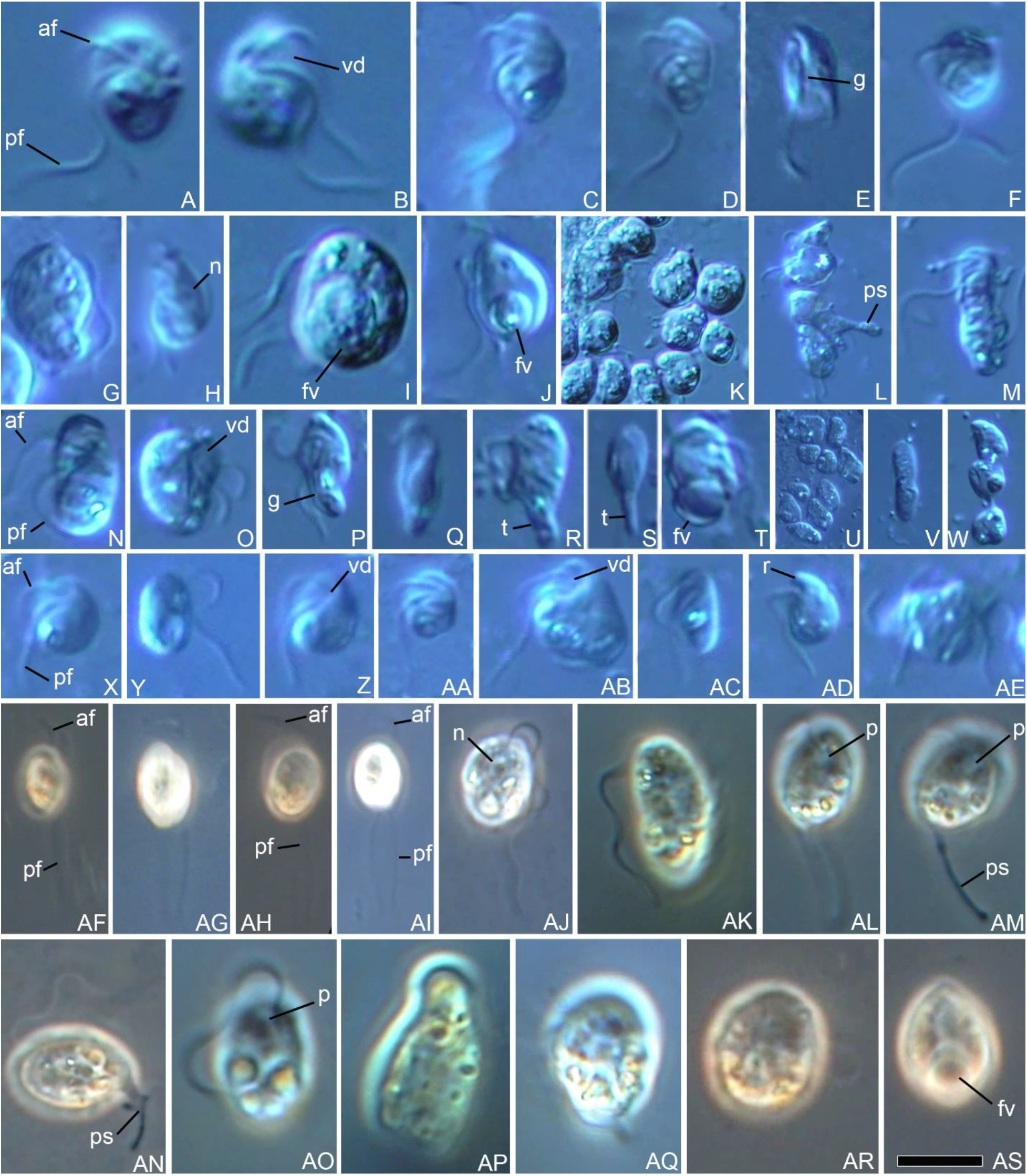
Morphology of novel phagoheterotrophic stramenopiles. **A, B.** *Develocanicus vyazemskyi,* general cell view with flagella (anterior flagellum [af] and posterior flagellum [pf]) and ventral depression [vd]. **C–M.** *Develocanicus komovi,* C–F – general cell view with flagella and ventral depression, shallow wide groove [g] is visible in (E), G – lying cell with posterior flagellum [pf] beating with a slow sinusoidal wave, H–J – cells with medial nucleus [n] (H) and large food vacuoles [fv] (I, J), K – cell aggregation, L – aggregated cells with pseudopodia [ps], M – transverse binary fission. **N–W.** *Develocauda condao,* N–P – general cell view with two flagella and ventral depression, Q – rod-shaped cell, R,S – cells with pointed ‘tail-like’ [t] posterior end, T – cells with large food vacuole, U – cell aggregation, V – transverse binary fission, W - partially fused cells. **X–AE.** *Сubaremonas variflagellatum,* X–AA – general cell view with flagella, AB – cell with conspicuous ventral depression, AC, AD – starving cells with small rostrum [r] (AD), AE – division into 4 cells. **AF, AG, AJ – AS.** *Feodosia pseudopoda,* AF, AG *–* typical fast swimming cell with two flagella, AJ, AK, AO – lying cells with sinusoid shaped flagella, AL–AN – cells with pseudopodia and anterior pit [p] (AL, AM), AP–AR – metabolic cells, AS – cell with large food vacuole. **AH.** *Koktebelia satura*, typical fast swimming cell with two flagella. **AI.** *Pirsonia chemainus*, typical fast swimming cell with two flagella. **Scale bar**: A, B, R, AK, AP, AQ, AS – 8 μm; C–H, J, N, P,Q, X–AA, AC, AD – 7 μm; I, O, T, AB – 5 μm; K, L, V, AF–AI – 15 μm; M, S, AE, AJ, AL–AO, AR – 10 μm; U – 25 μm; W – 20 μm.

The medial nucleus is located closer to the dorsal side of the cell (Fig. 4H). A large digestive vacuole is situated at the posterior part of the cell (Fig. 4I, J). As it is digested, the posterior end of the cell becomes thinner. The cells can form aggregations and attach to each other (Fig. 4K), sometimes form pseudopodia (Fig. 4L). Transverse binary fission (Fig. 4M).

#### Develocauda condao (Fig. 4N–W)

Free-swimming eukaryovorous heterokont flagellates (Colp-29c). The cells are slightly flattened, usually elongated-oval, less often narrow-oval or almost rod-shaped (Fig. 4Q). The anterior end is more rounded, the posterior end of the cell can be pointed, forming a characteristic “tail” found in starving cells (Fig. 4R, S). Cell length 5.14 - 12 μm, width 2.8 - 5.42 μm typically ranging 7.14 x 4.28 μm in dimension. The caudal extension is about 4.57 x 1.42 µm in size.

The cells have two heterodynamic flagella of an almost equal length with a posterior flagellum compared to the cell body. Flagella emerge from a pronounced deep ventral depression (Fig. 4N, O), which almost extends to the dorsal side of the cell. Depression transforms into a shallow groove (Fig. 4P) spanning along the entire cell, in which the posterior flagellum can fit.

The cells swim very quickly without rotating along the longitudinal axis. The posterior flagellum is straight and directed backwards. The anterior flagellum is directed forward, beats actively, and is only slightly curved. Rarely, the cells lie at the bottom with both flagella directed backward while making a slow sinusoidal movement, or the posterior flagellum beating actively.

The aggregated (Fig. 4U), partially fused cells (Fig. 4W) that form clusters were observed in culture. The medial nucleus is located closer to the dorsal side of the cell. Sated cells do not have a tail; at the posterior end of their cells there is a large digestive vacuole (Fig. 4T). Transverse binary fission (Fig. 4V).

#### Сubaremonas variflagellatum (Fig. 4X–AE)

Cells (clone Dev-1) are naked and solitary bacteriovores with the length 3.7 - 8 μm, the width 2.6 - 5.4 μm, and the typical dimension of 5.0 х 3.7 μm. The cell shape varies from elongated oval, oviform to rounder form (Fig. 4X-AA). Typically, the shape is irregularly ovoid, with the convex dorsal side and the flatter ventral side. The shape and size vary depending on feeding conditions. Starving cells have a small rostrum at the anterior end (Fig. 4AD). Cells are larger before division.

The cells possess two heterodynamic flagella of unequal length, emerging from a conspicuous ventral depression (Fig. 4Z, AB). Ventral depression starts from the anterior tip and continues ventrally to the middle of the cell. The anterior flagellum is approximately equal to the cell length or slightly longer while the posterior flagellum is 1.5 - 1.8 times longer than the cell. Digestive vacuole is situated at the cell posterior. An observed cell division produces two or four cells (Fig. 4AE).

In culture condition, the cells predominantly lie at the bottom unattached with both flagella directed backward. The posterior flagellum runs along the ventral surface of the cell and beats rapidly with sinusoidal pattern to draw water through the depression. The anterior flagellum is hook-shaped and sweeps slowly down behind the posterior flagellum.

Although less common, when the cells swim, the curved anterior flagellum actively beats, pulling the cell forward. It is almost invisible due to its fast beating. The posterior flagellum extends behind the cell and is likely used as a rudder. The cells swim quickly, only occasionally rotating about the axis of motion. Cells can sharply change the direction of movement.

Pirsoniales Cavalier-Smith 1998 emend. 2006

#### Feodosia pseudopoda (Fig. 4AF, AG, AJ–AS), Koktebelia satura (Fig. 4AH), and Pirsonia chemainus (Fig. 4AI)

Free-swimming naked, solitary and eukaryovorous heterokont flagellates. Cells are shaped as a flattened oval, with slightly pointed ends with the size 10.5 - 14 μm in length, 6 - 9.1 μm in width, and typically having the dimension of 12 x 8.2 μm. The flagellated stages of three studied Pirsoniales were almost morphologically identical except *Feodosia pseudopoda* (Chromo-2) which possesses small notch at the anterior part of the cell (Fig. 4AF, AG). Rarely, *F. pseudopoda* can produce pseudopodia (Fig. 4AM, AN), which are up to 10 μm long and sometimes branched.

Two long heterodynamic flagella originate from the pit located in the anterio-medial part of the cell (Fig. 4AL, AM, AO). The length of the anterior flagellum is as long as the cell, while the posterior one is 2.5 times longer.

The cells swim fast in a straight line, without rotating along the longitudinal axis. The anterior flagellum is directed anteriorly, always bent towards the ventral surface. The posterior flagellum propels the cell and beats at a high speed, which can be seen as multiple posterior flagella (Fig. 4AI). In stationary cells, the flagella take the form of a sinusoid (Fig. 4AJ, AK).

The nucleus is located in the middle of the cell (Fig. 4AJ). The cytoplasm contains many refractive granules as observed in previously described *Pirsonia* species (Schweikert and Schnepf, 1997). Non-flagellated cells were also observed with slightly amoeboid and round shape (Fig. 4AP–AR). The satiated cells have a large digestive vacuole at the posterior end (Fig. 4AS). The eukaryovory of the biflagellates seems to be facultative as the most did not actively pursue the prey but only *Koktebelia satura* (clone Chromo-1) consumed all the prey cells in culture.

## 4. Discussion

### 4.1 Monophyly and Phylogenetic Position of the Bigyromonadea

Of the known subdivisions of stramenopiles, the Bigyromonadea stand out for their lack of data and contentious position in the tree (even the newly discovered *P. tardus* is represented by transcriptomic data and consistency branches at the base of the tree). From the five recent phylogenomic analyses of stramenopiles, only three included a single bigyromonada representative (*D. elegans* JAMSTEC), none tested the monophyly of the group, and they recovered inconsistent positions. Using transcriptomes of seven new species belonging to the Bigyromonadea representing both the Developea and Pirsoniales subgroups, we tested the monophyly of the group and its position relative to other stramenopiles.

Previously, only SSU rRNA phylogenies could be used to test the monophyly of the Bigyromonadea, and such analyses consistently failed to support the monophyly, typically showing Developea with oomycetes and Pirsoniales with ochrophytes (Aleoshin et al., 2016; Kühn et al., 2004; Weiler et al., 2020). In contrast, phylogenomic data consistently and strongly supports the monophyly of these two groups, and shows each to include multiple distinct genera.

The position of Bigyromonadea within stramenopiles as a whole is also contentious, with some analyses showing the previously available transcriptome from *D. elegans* branching with oomycetes (Noguchi et al., 2016; Thakur et al., 2019), and based on internode consistency analyses (Kobert et al., 2016; Leonard et al., 2018, with ochrophytes). This discrepancy is not entirely eliminated by the addition of new taxa, since ML phylogenomic trees with the expanded representation recovered monophyly of the bigyromonada and oomycetes with robust support, but Bayesian analyses show a monophyly of bigyromonada+ochrophytes, and AU tests rejected most but not all topologies with this relationship (Table 1; Table S2).

The discrepancy between the ML and Bayesian analyses may be due to two groups (Chrysista and Bigyromonadea) that do not fit the same model for tree reconstruction. Although it is not the aim of this study to resolve the phylogeny of ochrophytes, further examination of ochrophyte phylogeny, may reveal whether the discrepancy stems from the unreconciled model used in the two groups, the different data processing approaches used, or insufficient data in one or both groups.

These results change how we interpret these lineages and their biological characteristics within the wider evolution of stramenopiles. For example, the phylogenetic position of Pirsoniales inferred from ribosomal genes showed they share a recent common ancestor with the ochrophytes, which naturally affected the interpretation of the ancestral state of ochrophytes and the role of phagoheterotrophy in their evolution (Aleoshin et al., 2016; Shiratori et al., 2017).

However, the phylogenomic tree points instead to a phagoheterotrophic origin of the Pseudofungi. Parallels between this and recent suggestions on the origin of fungi are noteworthy, since *Paraphelidium tribonemae*, a phagoheterotrophic parasite belonging to phylum Aphelida, has recently been found to be sister to the osmotrophic “core” fungi by phylogenomics (Torruella et al., 2018). Close similarities in metabolism and a phagotrophy-related proteome profile of *P. tribonemae* and the osmotrophic “core” fungi suggested the “core” fungi have evolved from a phagoheterotrophic aphelid-like ancestor. Further information on the metabolism and feeding mechanisms of the new species should shed light on whether the origins of fungi and pseudofungi have more parallels and on the possible phagoheterotrophic ancestral state of Gyrista more widely.

Of course, this is also dependent on conclusively determining the position of Bigyromonadea. Substantial advances in phylogenetic methods have been made, but challenges stemming from systematic errors, compositional bias, or long branch attraction, incomplete or contaminated data, and models that do not account for heterotachy in large datasets (Delsuc et al., 2005; Kapli et al., 2020; Zhou et al., 2007) remain. Similarly, advances in single-cell sequencing have vastly increased the taxonomic scope of phylogenomics, but the severely limited starting material and the fact that they are by definition a snapshot of gene expression in one cell remain important hurdles. Here, the removal of fast-evolving sites (Fig. 2), species (Fig. S3), extensive AU test (Table 1;Table S2;Fig. S2) and two different data processing approaches collectively tip the scale in favour of the monophyly of bigyromonada and oomycetes over the alternative position of bigyromonada with ochrophytes. However, the conflicting results of Bayesian inferences show that the lack of a robust phylogenomic tree was not just due to lack of taxonomic diversity. Continued sampling efforts in phagoheterotrophic stramenopiles will expand the phylogenetic diversity of the Bigyromonadea (and environmental SSU rRNA data already show there are more new taxa to be found), but other advances in data generation and analyses will also be required.

### 4.2 Morphology, evolutionary implications, and taxonomic description of the novel phagoheterotrophic Bigyromonadea

#### 4.2.1 Newly observed morphological and behavioural features in bigyromonads: cell-aggregation to fusion, pseudopod-formation, and facultative phagotrophy in motile zoospores

Before we compare morphological features, we need to clarify that the JAMSTEC strain of *Developayella elegans* has been mis-named and is a distinct species in a different genus.

According to the SSU rRNA gene tree (Fig. 3), the originally described *D. elegans* U37107 (Tong, 1995), is only distantly related to *D. elegans* JAMSTEC, and placed in distinct sub-clade of Developea where it is most closely related to *C. variflagellatum*. Renaming *D. elegans* JAMSTEC will be necessary in the future: its close relatedness to *Cubaremonas* is sufficient to say it is mis-named, but rectifying this should take into account morphological information, which is currently unavailable. Overall, however, the novel developeans have similar morphological traits as previously described species. For example, *C. variflagellatum* falls in the same sub-clade as *Mediocremonas mediterraneus* (Del Campo et al., 2013; Weiler et al., 2020) (Fig. 3), and both have similar morphology. *C. variflagellatum* is slightly larger, but measurements for *M. mediterraneus* (2.0 - 4.0 μm in length and 1.2 - 3.7 μm in width) were most likely based on scanning electron microscopy (SEM) images and cells tend to shrink in SEM fixatives (Weiler et al., 2020). The cell size, flagella length and swimming movement of *C. variflagellatum* exhibited close similarity to *D. elegans* U37107, which was named after its characteristic developpé movement of the anterior flagellum during stationary feeding (Tong, 1995). However, no thread-like substances were observed, which *D. elegans* uses to attach to substrate.

The remaining novel Developea species, *Develocanicus vyazemskyi, D. komovi*, and *Develocauda condao,* differed from *D. elegans* JAMSTEC and *C. variflagellatum* by having a proportionately longer posterior flagellum, forward propulsion without rotating its axis, a eukaryovorous diet (like *Develorapax marinus* (Aleoshin et al., 2016)), and the presence of a “tail” in *D. condao*. Notably, the ability of the cells to form aggregates (Fig. 4K, U), pseudopodia (Fig. 4L), and to undergo partial cell fusion (Fig. 4W) has not been reported in this clade previously. The above-mentioned differences between *D. vyazemskyi D. komovi*, *Develocauda condao*, and *C. variflagellatum* are also phylogenetically reflected in the division of these species into two sub-clades (Fig. 1; Fig. 3).

The three novel Pirsoniales, *Feodosia pseudopoda, Koktebelia satura*, and *Pirsonia chemainus,* described here as *nomen provisorium,* most likely represent a motile zoospore stage of unknown algal parasites. The novel Pirsoniales species did not actively pursue the provided prey and only partially consumed their prey (except *K. satura* which consumed all the prey provided), all the cultures died after a few months to one year of cultivation. Although there has been extensive description of auxosome and trophosome formation during the parasitic stage of known Pirsoniales (Schnepf et al., 1990; Schweikert and Schnepf, 1997), the ability of motile zoospores to acquire effective eukaryovory has not been described so far. The observed eukaryovory of the zoospore-like Pirsoniales is likely facultative, as the cells were cultured without potential hosts and the cells with larger food vacuoles became non-flagellated and rounded, a structure akin to an auxosome. However, further culture experimentations with their natural hosts are required to verify their ability to form parasitic auxosomes and trophosomes from motile phagotrophic zoospores.

We postulate that the facultative eukaryovory at the motile zoospore stage provides a significantly increased survival rate and thus extension of the motile stage during their dispersal until a suitable host is found. This ability can be particularly advantageous before the onset of seasonal algal bloom, where the zoospores can efficiently infect multiple hosts without resource competition. Therefore, the sustained survival of the zoospores via facultative eukaryovory could be an important factor leading to the evolutionary success of Pirsoniales parasites.

*Feodosia pseudopoda* differed from rest of the Pirsoniales studied here by an anterior notch (Fig. AF, AG) and rare occurrences of pseudopodia (Fig. AM-AO). Interestingly, the two characteristics have been reported in *Pseudopirsonia mucosa*, a cercomonad rhizarian (Kühn et al., 2004), which had been mis-assigned as *Pirsonia* due to the similarities in their parasitic life cycles. In starving and immobile zoospores of *Pirsonia puntigerae*, filopodium-like processes (Schweikert and Schnepf, 1997) have been described however, pseudopodia in motile zoospores of Pirsoniales have not been observed previously.

The presence of pseudopodia, and the ability to form aggregated cells in the newly described sub-clade I of Developea and previously reported publications of Pirsoniales may indicate synapomorphic traits of Bigyromonadea. It will be important for future studies to compare ultrastructure and genes putatively associated with cell-aggregation or fusion among the species of bigyromonada, thus potentially addressing the evolution of an osmotrophic nutritional strategy in stramenopiles.

#### 4.2.2 Similarities among Oomycetes motile zoospores, Labyrinthulomycetes, and Bigyromonadea

Morphologically, the novel Developea species have similar features as motile zoospores of previously studied oomycetes, such as the general cell dimension, the proportion between anterior and posterior flagellum, and two laterally oriented flagella (with a tinsellate anterior flagellum) emerging from a ventral groove (Dick, 2000), which resembles the ventral depression observed in the novel species. Behaviourally, the swimming pattern (e.g., direction of flagella, sinusoid form) is comparable (Hickman, 1970; Ho and Hickman, 1967). Another striking similarity between the two groups is their ability to self-aggregate, which is observed in oomycete zoospores as a distinct form of self-aggregation compared to aggregation towards host-plant tissues (Bassani et al., 2020; Ko and Chase, 1973). The mechanism underlying self-aggregation in oomycetes has not been fully resolved, however recent studies suggest that a combination of chemotaxis (Bassani et al., 2020; Judelson and Blanco, 2005; Zheng and Mackrill, 2016) and bioconvection (Savory et al., 2014), is involved in the process. The exact role of the self-aggregation in oomycete pathogenesis is still unclear, however the fact that a similar observation was made in its sister-clade, the Bigyromonadea, indicates that self-aggregation may have been present in the ancestor of Pseudofungi, before the osmotrophic parasitism of oomycetes evolved. Cell aggregation is also observed in *Sorodiplophrys* (Dykstra et al., 1975), a species belonging to another osmotrophic group of stramenopiles, the labyrinthulomycetes. Cell aggregation has convergently evolved multiple times across many other supergroups (Parfrey and Lahr, 2013), such as Amoebozoa (Du et al., 2015), Rhizaria (Brown et al., 2012), and ciliates (Sugimoto and Endoh, 2006), and whether cell aggregation within stramenopiles arose convergently or divergently should be further investigated.

As mentioned previously, some species described in this study formed pseudopodia (Fig. 4L,4AM,4AN) and partially fused cells (Fig. 4W) resembling amoeboid forms. Interestingly, labyrinthulomycetes also form filose pseudopodia (Gomaa et al., 2013) akin to pseudopodia observed in this study (Fig. 4AM, AN). These are found in Amphitremidae, during an amoeboid stage of *Diplophrys* (Anderson and Cavalier-Smith, 2012), and other labyrinthulids (Raghukumar, 1992), implying this trait either evolved convergently or was present earlier than the divergence of Pseudofungi.

Another notable similarity between oomycetes and the novel bigyromonada is their potential marine origin, as all known bigyromonads are exclusively marine. Molecular clock analyses indicate the Silurian period as the time of oomycete origins (Matari and Blair, 2014), while the earliest fossil evidence points to the Devonian period (Krings et al., 2011). The fossil evidence of the early diverging genera have shown them to be marine parasites of seaweed, or of crustaceans based on molecular studies (Beakes and Sekimoto, 2009), both suggesting a marine origin of oomycetes as a facultative parasitic osmotroph (Beakes et al., 2014, 2012; Beakes and Thines, 2017). The origin and evolution of major stramenopile subgroups is coming into sharper focus with the increase in phylogenomic data from diverse species. The new taxa described here, together with future descriptions of the still-substantial diversity of bigyromonada that has not been well-characterized, can potentially shed more light on this and the origins of oomycetes in particular. We propose that the ancestor of oomycetes was a phagoheterotrophic amoeboid, as postulated in the evolution of true fungi (Zmitrovich, 2018), and that this transition might be better understood through a detailed functional examination of the novel species. Just as the highly successful analyses of choanoflagellates and unicellular opisthokonts changed our understanding of the origin of animals (Chow et al., 2019; Sebé-Pedrós et al., 2013), a similar analysis of the distribution of genes involved in Pseudofungi cell-aggregation or pseudopodia formation across the diversity of bigyromonads could be a future direction to understand the evolution of these unique phagoheterotrophs and oomycetes.

## TAXONOMIC SUMMARY

Taxonomy: Eukaryota; SAR Burki et al. 2008, emend. Adl et al. 2012; Stramenopiles Patterson 1989, emend. Adl et al. 2005; Gyrista Cavalier-Smith 1998; Bigyromonadea Cavalier-Smith, T. 1998; Developea Karpov et Aleoshin 2016 *Сubaremonas* n. gen. Tikhonenkov, Cho, and Keeling Diagnosis: naked and solitary bacteriovorous protist. Cell shape is irregularly ovoid, with the convex dorsal side and the flatter ventral side. Cells possess two heterodynamic flagella emerging from a conspicuous ventral depression, which starts from the anterior end and continues ventrally to the middle of the cell. In culture condition, the cells predominantly lie at the bottom unattached with both flagella directed backward.

Etymology: from lat. cubare – to lie, to be lying down and monas (lat.) – unicellular organism.

Zoobank Registration. urn:lsid:zoobank.org:act:xxxxx

Type species. *Сubaremonas variflagellatum*

*Сubaremonas variflagellatum* n. sp. Tikhonenkov, Cho, and Keeling Diagnosis: cells length 3.7 - 8 μm, cell width 2.6 - 5.4 μm. Flagella of unequal length, the anterior one is approximately equal to the cell length while the posterior flagellum is 1.5 - 1.8 times longer than the cell. At lying cells, posterior flagellum runs along the ventral surface of the cell and beats rapidly with sinusoidal pattern to draw water through the depression. The anterior flagellum is hook-shaped and sweeps slowly down behind the posterior flagellum. Starving cells have a small rostrum at the anterior end. Digestive vacuole is situated at the cell posterior. An observed cell division produces two or four cells.

Type Figure: Fig. 4X illustrates a live cell of strain Dev-1.

Gene sequence: The SSU rRNA gene sequence has the GenBank Accession Number XXXXX. Type locality: water column of Strait of Georgia, British Columbia, Canada

Etymology: the species name means “unequal flagella”, lat.

Zoobank Registration: urn:lsid:zoobank.org:act:XXXXXXXX

*Develocanicus* n. gen. Tikhonenkov, Cho, Mylnikov, and Keeling

Diagnosis: Free-swimming naked eukaryovorous heterokont flagellates with two non- acronematic heterodynamic flagella of unequal lengths. The shape of the cell is irregularly flattened ellipse, where the dorsal side is more convex, and the ventral side is flatter. Flagella emerge from a prominent ventral depression which passes into a shallow wide groove along the entire length of the cell.

Etymology: from développé (fr.) – characteristic ballet movement and volcanicus (lat.) (found near volcanos in Kanary island and Crimea).

Zoobank Registration. urn:lsid:zoobank.org:act:xxxxx

Type species. *Develocanicus komovi*

*Develocanicus komovi* n. sp. Tikhonenkov, Cho, Mylnikov, and Keeling

Diagnosis: cell length 5.4 - 10 μm, cell width 3.8 - 7.4 μm. The posterior flagellum is two times longer than the cell, the anterior flagellum is approximately 1 - 1.5 times longer. Cells swim without rotation. At that, posterior flagellum is directed backward and straight, running along the ventral cell of the cell. Anterior flagellum beats rapidly and is directed forward while slightly curved. Medial nucleus is located closer to the dorsal side of the cell. Large digestive vacuole is situated at the posterior part of the cell. Cells can form pseudopodia and aggregations and attach to each other. Transverse binary fission.

Type Figure: Fig. 4C illustrates a live cell of strain Colp-23.

Gene sequence: The SSU rRNA gene sequence has the GenBank Accession Number XXXXX.

Type locality: black volcanic sand on the littoral of Maria Jimenez Beach (Playa Maria Jiménez), Puerto de la Cruz, Tenerife, Spain

Etymology: named after Prof., Dr. Viktor T. Komov, Russian ecotoxicologist, who carried out fieldwork and collect samples, where new species was discovered.

Zoobank Registration: urn:lsid:zoobank.org:act:XXXXXXXX

*Develocanicus vyazemskyi* n. sp. Tikhonenkov, Cho, Mylnikov, and Keeling

Diagnosis: cell 7.4 - 12.5 μm long, 4.8 - 9.2 μm wide. The posterior flagellum is two times longer than the cell, the anterior flagellum is approximately 1 - 1.5 times longer. Cells swim without rotation. At that, posterior flagellum is directed backward and straight, running along the ventral cell of the cell. Anterior flagellum beats rapidly and is directed forward while slightly curved. In non-motile cells, both flagella are directed backward, beating in a slow sinusoidal wave. Medial nucleus is located closer to the dorsal side of the cell. Large digestive vacuole is situated at the posterior part of the cell. Transverse binary fission.

Type Figure: Fig. 4A illustrates a live cell of strain Colp-30.

Gene sequence: The SSU rRNA gene sequence has the GenBank Accession Number XXXXX.

Type locality: near shore sediments on the littoral near T.I. Vyazemsky Karadag Scientific Station, Crimea

Etymology: named after Dr. T.I. Vyazemsky, founder and first director of Karadag Scientific Station, Crimea

Zoobank Registration: urn:lsid:zoobank.org:act:XXXXXXXX

*Develocauda* n. gen. Tikhonenkov, Cho, and Keeling

Diagnosis: Free-swimming eukaryovorous heterokont flagellates with slightly flattened elongated-oval cells and two heterodynamic flagella. The anterior end is more rounded, the posterior end of the cell can be pointed, forming a characteristic “tail” in starving cells. Flagella emerge from a pronounced deep ventral depression, which almost extends to the dorsal side of the cell. Depression transforms into a shallow groove spanning along the entire cell, in which the posterior flagellum can fit.

Etymology: from développé (fr.) – characteristic ballet movement and cauda (lat.) – tail.

Zoobank Registration. urn:lsid:zoobank.org:act:xxxxx Type species. *Develocauda condao*

*Develocauda condao* n. sp. Tikhonenkov, Cho, and Keeling

Cell length 5.14 - 12 μm, width 2.8 - 5.42 μm. The caudal extension is about 4.57 x 1.42 µm in size. Flagella of almost equal length. The cells swim very quickly without rotating along the longitudinal axis. The posterior flagellum is straight and directed backwards. The anterior flagellum is directed forward, beats actively, and is only slightly curved. Cells can be partially fused and aggregated. Medial nucleus is located closer to the dorsal side of the cell. Transverse binary fission.

Type Figure: Fig. 4N illustrates a live cell of strain Colp-29.

Gene sequence: The SSU rRNA gene sequence has the GenBank Accession Number XXXXX.

Type locality: near shore sediments on the littoral of north-east part of Con Dao Island, South Vietnam

Etymology: named after Con Dao Island, South Vietnam, where species was discovered.

Zoobank Registration: urn:lsid:zoobank.org:act:XXXXXXXX

Pirsoniales Cavalier-Smith 1998, emend. 2006

Studied pirsoniales most likely represent a motile zoospore stages of unknown algal parasites. Since data on the stage of the parasitic trophonts (auxosome and a trophosome) are not available, it is premature to formulate taxonomic diagnoses. But we provide provisional names (nom. prov.) which can be used for future research.

*Pirsonia chemainus* nom. prov. Tikhonenkov, Cho, and Keeling

Etymology: species epithet is after the Stz’uminus First Nation traditional territory (Strait of Georgia area) claimed by the Chemainus First Nation

Type locality: water column of the Strait of Georgia, British Columbia, Canada

Gene sequence: The SSU rRNA gene sequence has the GenBank Accession Number XXXXX.

*Koktebelia satura* nom. prov. Tikhonenkov, Cho, and Keeling

Etymology: genus epithet reflects the place of finding, Koktebel bay, Crimea; species epithet – from satur (lat.), well-fed.

Type locality: near shore sediments on the littoral near T.I. Vyazemsky Karadag Scientific Station, Crimea

Gene sequence: The SSU rRNA gene sequence has the GenBank Accession Number XXXXX.

*Feodosia pseudopoda* nom. prov. Tikhonenkov, Cho, and Keeling

Etymology: genus epithet reflects the place of finding, the settlement Beregovoye, Feodosiya, Crimea; species epithet reflects the ability to produce pseudopodia.

Type locality: near shore sand on the littoral of the beach in the settlement Beregovoye, Feodosiya, Crimea

Gene sequence: The SSU rRNA gene sequence has the GenBank Accession Number XXXXX.

## Acknowledgements

We thank Dr. Viktor Komov (IBIW RAS), Larysa Pakhomova (UBC), Dr. Evgeny Gusev (IPP RAS) for helping with the sample collection in the Canary Islands, Strait of Georgia (British Columbia), and Vietnam as well as Dmitry Zagumyonnyi (IBIW RAS) for the help with Pirsoniales photography. Field work in Vietnam is part of the project ‘Ecolan 3.2’ of the Russian-Vietnam Tropical Centre. This work was supported by the Russian Science Foundation grant no. 18-14-00239, https://rscf.ru/project/18-14-00239/ (cell isolation and culturing, microscopy, SSU rRNA sequencing, and analyses), NSERC Grant 2014- 03994 to PJK (sequencing and bioinformatics), and NSERC Postgraduate Scholarship-Doctoral (PGSD) and University of British Columbia Botany Four-Year Fellowship to AC.

