## Supplemental Table 1 for "Monophyly of Diverse Bigyromonadea and their Impact on Phylogenomic Relationships Within Stramenopiles"

**Supplemental Material**

**Table S1**. List of 27 recently published Stramenopiles taxa and the corresponding genome or transcriptome data included in this study. 'Peptide reads' were extracted from an annotated genome sequences and publicly available Marine Microbial Eukaryote Transcriptome Sequencing Project (MMETSP) website (https://www.imicrobe.us/#/projects/104). 'Transcripts' were extracted from an unannotated genome sequences, and protein sequences were predicted as described in Method.

| Group | Sample ID | Type | Project | SRA Run | GenBank | Strain |
|---|---|---|---|---|---|---|
| Ochrophytes | *Rhizosolenia setigera* (MMETSP0789) | Peptide reads | PRJNA248394 | SRR1296707 | | CCMP 1694 |
| | *Synchroma pusilum* (MMETSP1452) | Peptide reads | PRJNA248394 | SRR1300531 | | CCMP 3072 |
| | *Aureococcus anophagefferens* | Peptide reads | PRJNA13500 | | GCA_000186865.1 | CCMP 1984 |
| Oomycetes | *Phytophthora parasitica* | Peptide reads | PRJNA73155 | | GCA_000247585.2 | INRA-310 |
| | *Plasmopara halstedii* | Peptide reads | PRJEB6932 | | GCA_900000015.1 | |
| | *Hyaloperonospora arabidopsidis* | Peptide reads | PRJNA298674 | | GCA_001414525.1 | Noks1 |
| | *Nothophytophthora sp.* | Peptide reads | PRJNA328215 | | GCA_001712635.2 | Chile5 |
| | *Pythium ultimum* | Peptide reads | PRJNA36503 | | GCA_000143045.1 | DAOM BR144 |
| | *Pythium brassicum* | Peptide reads | PRJNA498716 | | GCA_008271595.1 | P1 |
| | *Albugo candida* | Peptide reads | PRJNA291031 | | GCA_001306755.1 | Ac 7v |
| | *Saprolegnia diclina* | Peptide reads | PRJNA86859 | | GCA_000281045.1 | VS20 |
| | *Achlya hypogyna* | Peptide reads | PRJNA169234 | | GCA_002081595.1 | ATCC 48635 |
| | *Thraustotheca clavata* | Peptide reads | PRJNA169235 | | GCA_002081575.1 | ATCC 34112 |
| | *Aphanomyces astaci* | Peptide reads | PRJNA187372 | | GCA_000520075.1 | APO3 |

| | | | | | | |
|---|---|---|---|---|---|---|
| | *Hyphochytrium catenoides* | Transcripts | PRJEB13950 | | GCA_900088475.1 | |
| Developea | *Developayella elegans* | Transcriptome | PRJDB4370 | DRR049556 | | CCAP:1917/1 |
| Sagenista | *Pseudophyllomitus vesiculosus* | Transcriptome | PRJDB8568 | DRR186658 | | NIES-4114 |
| | MAST4 | Peptide reads | PRJNA244411 | SRR1263007 | | dcp33 |
| | MAST4A2 | Transcripts | PRJEB6603 | | GCA_900128395.1 | TOSAG23-2 |
| | MAST4E | Transcripts | PRJEB6603 | | GCA_900128585.1 | TOSAG23-3 |
| Oplozoa | *Wobblia lunata* | Transcriptome | PRJDB4369 | DRR049555 | | NIES-1015 |
| | *Blastocystis*_ST4 | Peptide reads | PRJNA257240 | | GCA_000743755.1 | WR1 |
| | MAST3 | Transcripts | PRJEB6603 | ERR1198953 | | TOSAG41-2 |
| | MAST3F | Transcripts | PRJEB6603 | | GCA_900128565.1 | TOSAG23-6 |
| | *Cafeteria roenbergensis* | Peptide reads | PRJNA552725 | | GCA_008330645.1 | BVI |
| | *Bicosoecid* sp. (MMETSP0115) | Peptide reads | PRJNA231566 | SRR1294380 | | |
| | *Cantina marsupialis* | Transcriptome | PRJDB3523 | DRR030401 | | YPF1205 |
| Platysulcea | *Platysulcus tardus* | Transcriptome | PRJDB8466 | DRR186656 | | NIES-3720 |
