## Supplemental Figure 1 for "Monophyly of Diverse Bigyromonadea and their Impact on Phylogenomic Relationships Within Stramenopiles"

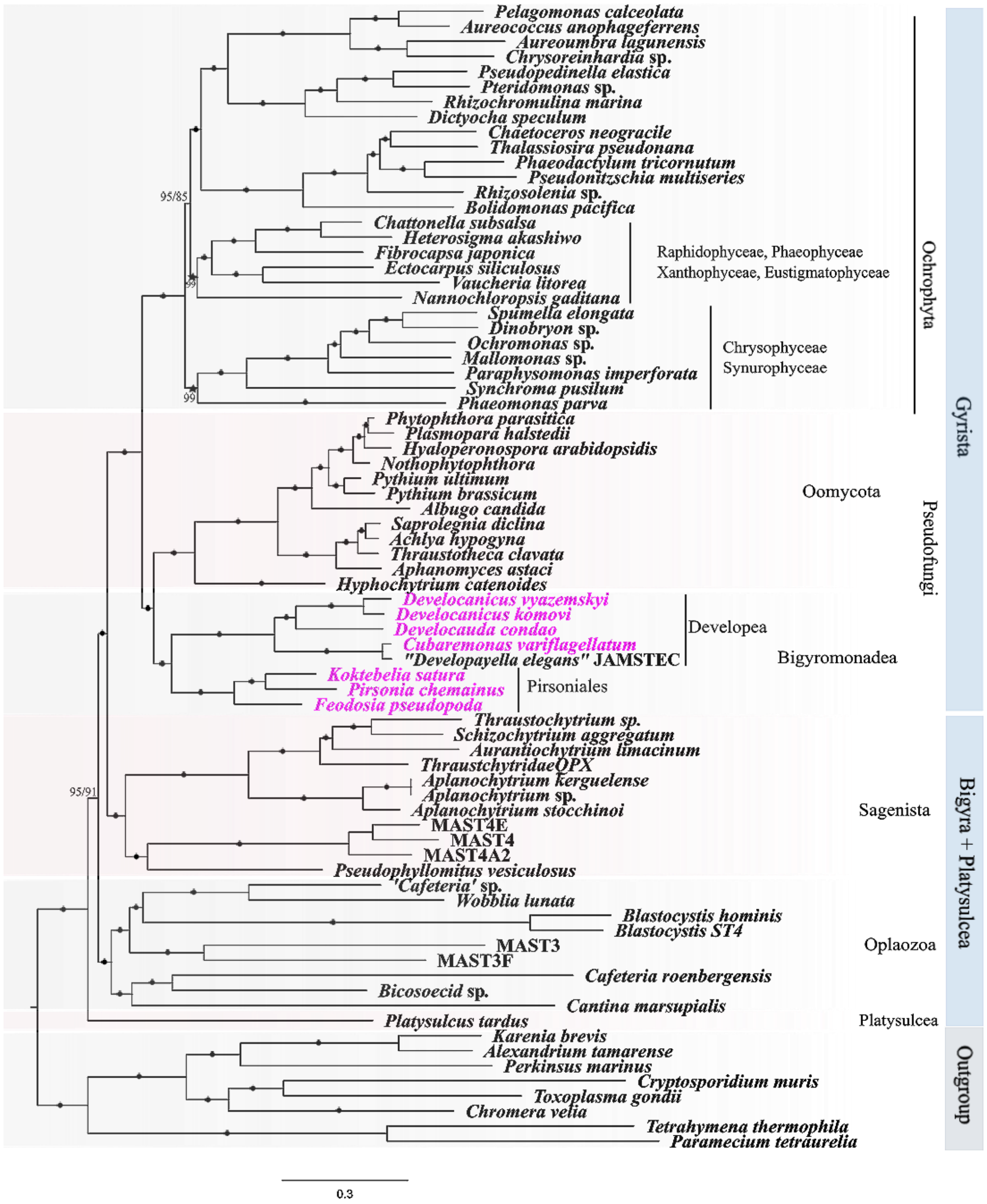


**Figure S1.** Multi-gene phylogenomic tree of stramenopiles with the seven new transcriptomes (pink) added to Gyrista, consisting of the concatenated alignments of 247 aligned gene-sets. The tree was reconstructed using the Maximum-likelihood (ML) analysis, under the site-heterogenous model, LG+C60+F+G4+PMSF implemented in IQ-Tree. The tree topology is based on the tree reconstructed on the dataset process with approach 2. Branch support was calculated separately using non-parametric PMSF 100 standard bootstrap (STB) from the dataset processed with two approaches. Branches with ≥99% STB for both approaches are marked with black bullets while others are labelled as “Approach 1 STB/Approach 2 STB”. The topology of the trees generated from the two approaches were the same except the positions of Raphidophyceae + Phaeophyceae + Xanthophyceae + Eustigmatophacea and Chrysophyceae + Synurophyceae, which were swapped in the tree reconstructed based on the dataset processed using approach 1; denoted by star symbols (Fig. 1).
