## Supplemental Figure 2 for "Monophyly of Diverse Bigyromonadea and their Impact on Phylogenomic Relationships Within Stramenopiles"

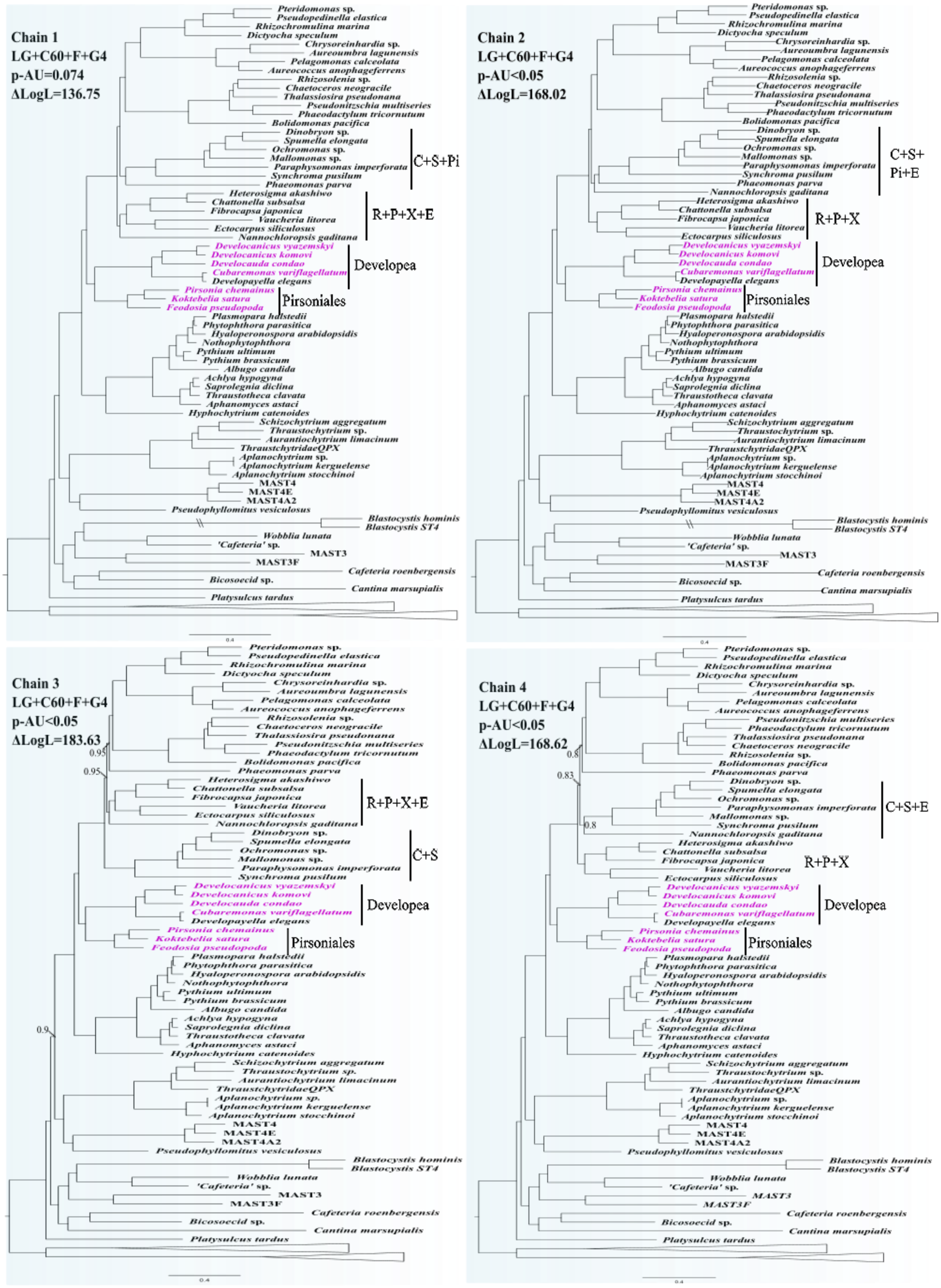
Chain 1
LG+C60+F+G4
p-AU=0.074
ΔLogL=136.75
Pteridomonas sp.
Pseudopedinella elastica
Rhizochromulina marina
Dictyocha speculum
Chrysoreinhardia sp.
Aureoumbra lagunensis
Pelagomonas calceolata
Aureococcus anophageferrens
Rhizosolenia sp.
Chaetoceros neogracile
Thalassiosira pseudonana
Pseudonitzschia multiseries
Phaeodactylum tricornutum
Bolidomonas pacifica
Dinobryon sp.
Spumella elongata
Ochromonas sp.
Mallomonas sp.
Paraphysomonas imperforata
Synchroma pusilum
Phaeomonas parva
C+S+Pi
Heterosigma akashiwo
Chattonella subsalsa
Fibrocapsa japonica
Vaucheria litorea
Ectocarpus siliculosus
Nannochloropsis gaditana
R+P+X+E
Develocanicus vyazemskyi
Develocanicus komovi
Develocauda condao
Cubaremonas variflagellatum
Developayella elegans
Developea
Pirsonia chemainus
Koktebelia satura
Feodosia pseudopoda
Pirsoniales
Plasmopara halstedii
Phytophthora parasitica
Hyaloperonospora arabidopsidis
Nothophytophthora
Pythium ultimum
Pythium brassicum
Albugo candida
Achlya hypogyna
Saprolegnia diclina
Thraustotheca clavata
Aphanomyces astaci
Hyphochytrium catenoides
Schizochytrium aggregatum
Thraustochytrium sp.
Aurantiochytrium limacinum
ThraustchytridaeQPX
Aplanochytrium sp.
Aplanochytrium kerguelense
Aplanochytrium stocchinoi
MAST4
MAST4E
MAST4A2
Pseudophyllomitus vesiculosus
Blastocystis hominis
Blastocystis ST4
Wobblia lunata
'Cafeteria' sp.
MAST3
MAST3F
Cafeteria roenbergensis
Bicosoecid sp.
Cantina marsupialis
Platysulcus tardus
0.4
Chain 2
LG+C60+F+G4
p-AU<0.05
ΔLogL=168.02
Pteridomonas sp.
Pseudopedinella elastica
Rhizochromulina marina
Dictyocha speculum
Chrysoreinhardia sp.
Aureoumbra lagunensis
Pelagomonas calceolata
Aureococcus anophageferrens
Rhizosolenia sp.
Chaetoceros neogracile
Thalassiosira pseudonana
Pseudonitzschia multiseries
Phaeodactylum tricornutum
Bolidomonas pacifica
Dinobryon sp.
Spumella elongata
Ochromonas sp.
Mallomonas sp.
Paraphysomonas imperforata
Synchroma pusilum
Phaeomonas parva
Nannochloropsis gaditana
C+S+
Pi+E
Heterosigma akashiwo
Chattonella subsalsa
Fibrocapsa japonica
Vaucheria litorea
Ectocarpus siliculosus
R+P+X
Develocanicus vyazemskyi
Develocanicus komovi
Develocauda condao
Cubaremonas variflagellatum
Developayella elegans
Developea
Pirsonia chemainus
Koktebelia satura
Feodosia pseudopoda
Pirsoniales
Plasmopara halstedii
Phytophthora parasitica
Hyaloperonospora arabidopsidis
Nothophytophthora
Pythium ultimum
Pythium brassicum
Albugo candida
Achlya hypogyna
Saprolegnia diclina
Thraustotheca clavata
Aphanomyces astaci
Hyphochytrium catenoides
Schizochytrium aggregatum
Thraustochytrium sp.
Aurantiochytrium limacinum
ThraustchytridaeQPX
Aplanochytrium sp.
Aplanochytrium kerguelense
Aplanochytrium stocchinoi
MAST4
MAST4E
MAST4A2
Pseudophyllomitus vesiculosus
Blastocystis hominis
Blastocystis ST4
Wobblia lunata
'Cafeteria' sp.
MAST3
MAST3F
Cafeteria roenbergensis
Bicosoecid sp.
Cantina marsupialis
Platysulcus tardus
0.4
Chain 3
LG+C60+F+G4
p-AU<0.05
ΔLogL=183.63
Pteridomonas sp.
Pseudopedinella elastica
Rhizochromulina marina
Dictyocha speculum
Chrysoreinhardia sp.
Aureoumbra lagunensis
Pelagomonas calceolata
Aureococcus anophageferrens
Rhizosolenia sp.
Chaetoceros neogracile
Thalassiosira pseudonana
Pseudonitzschia multiseries
Phaeodactylum tricornutum
0.95
Bolidomonas pacifica
0.95
Phaeomonas parva
Heterosigma akashiwo
Chattonella subsalsa
Fibrocapsa japonica
Vaucheria litorea
Ectocarpus siliculosus
Nannochloropsis gaditana
R+P+X+E
Dinobryon sp.
Spumella elongata
Ochromonas sp.
Mallomonas sp.
Paraphysomonas imperforata
Synchroma pusilum
C+S
Develocanicus vyazemskyi
Develocanicus komovi
Develocauda condao
Cubaremonas variflagellatum
Developayella elegans
Developea
Pirsonia chemainus
Koktebelia satura
Feodosia pseudopoda
Pirsoniales
Plasmopara halstedii
Phytophthora parasitica
Hyaloperonospora arabidopsidis
Nothophytophthora
Pythium ultimum
Pythium brassicum
Albugo candida
Achlya hypogyna
Saprolegnia diclina
Thraustotheca clavata
Aphanomyces astaci
0.9
Hyphochytrium catenoides
Schizochytrium aggregatum
Thraustochytrium sp.
Aurantiochytrium limacinum
ThraustchytridaeQPX
Aplanochytrium sp.
Aplanochytrium kerguelense
Aplanochytrium stocchinoi
MAST4
MAST4E
MAST4A2
Pseudophyllomitus vesiculosus
Blastocystis hominis
Blastocystis ST4
Wobblia lunata
'Cafeteria' sp.
MAST3
MAST3F
Cafeteria roenbergensis
Bicosoecid sp.
Cantina marsupialis
Platysulcus tardus
0.4
Chain 4
LG+C60+F+G4
p-AU<0.05
ΔLogL=168.62
Pteridomonas sp.
Pseudopedinella elastica
Rhizochromulina marina
Dictyocha speculum
Chrysoreinhardia sp.
Aureoumbra lagunensis
Pelagomonas calceolata
Aureococcus anophageferrens
Pseudonitzschia multiseries
Phaeodactylum tricornutum
Thalassiosira pseudonana
Chaetoceros neogracile
0.8
Rhizosolenia sp.
Bolidomonas pacifica
0.83
Phaeomonas parva
Dinobryon sp.
Spumella elongata
Ochromonas sp.
Paraphysomonas imperforata
Mallomonas sp.
0.8
Synchroma pusilum
Nannochloropsis gaditana
C+S+E
Heterosigma akashiwo
Chattonella subsalsa
Fibrocapsa japonica
Vaucheria litorea
Ectocarpus siliculosus
R+P+X
Develocanicus vyazemskyi
Develocanicus komovi
Develocauda condao
Cubaremonas variflagellatum
Developayella elegans
Developea
Pirsonia chemainus
Koktebelia satura
Feodosia pseudopoda
Pirsoniales
Plasmopara halstedii
Phytophthora parasitica
Hyaloperonospora arabidopsidis
Nothophytophthora
Pythium ultimum
Pythium brassicum
Albugo candida
Achlya hypogyna
Saprolegnia diclina
Thraustotheca clavata
Aphanomyces astaci
Hyphochytrium catenoides
Schizochytrium aggregatum
Thraustochytrium sp.
Aurantiochytrium limacinum
ThraustchytridaeQPX
Aplanochytrium sp.
Aplanochytrium kerguelense
Aplanochytrium stocchinoi
MAST4
MAST4E
MAST4A2
Pseudophyllomitus vesiculosus
Blastocystis hominis
Blastocystis ST4
Wobblia lunata
'Cafeteria' sp.
MAST3
MAST3F
Cafeteria roenbergensis
Bicosoecid sp.
Cantina marsupialis
Platysulcus tardus
0.4

**Figure S2.** Bayesian phylogenomic tree of stramenopiles of the seven new transcriptomes (pink) added to Gyrista. The tree was reconstructed based on the 247 gene-sets of 76 taxa processed with the approach 1 using PhyloBayes under the CAT+GTR+G4 model. No chains converged (maxdiff=1) and all chains have identical tree topologies except the sub-clades of ochrophytes as summarized as different combination of Raphidophyceae (R), Eustigmatophyceae (E), Chrysophyceae (C), Synurophyceae (S), Phaeophyceae (P), Pinguiophyceae (Pi), and Xanthophyceae (X). *Rhizosolenia* sp., also showed inconsistent topology across the chains. P-values were calculated using the approximately unbiased test (p-AU) with 10,000 RELL bootstrap replicates, implemented in IQ-TREE. The difference in maximum log likelihoods ($\Delta$LogL) of each tree was calculated by comparing to the maximum log likelihood of ML tree reconstructed under the LG+C60+F+G4. Except chain1, the topologies of the trees from chain 2-4 were rejected where their p-AU were less than 0.05, indicating confidence interval below 95%. Only the Bayesian posterior probabilities (PP) lower than 1 are marked in the figure. All other nodes have PP=1. The collapsed clades in the figures indicate outgroup (alveolates).
