## Supplemental Table 2 for "Monophyly of Diverse Bigyromonadea and their Impact on Phylogenomic Relationships Within Stramenopiles"

**Table S2**. Approximately unbiased (AU) test of constrained trees based on approach 2 dataset.

| Approach 2 (Prequal/Divvier, MAFFT G-INS-i, -gt 0.1) | | | |
|---|---|---|---|
| Constrained Tree | p-AU | logL | ΔlogL |
| Unconstrained ML tree | 0.602 | -4112709.551 | 0 |
| ML tree under LG+C60+F+G4+PMSF | 0.569 | -4112709.552 | 0.00035827 |
| ML tree under LG+C60+F+G4 | 0.543 | -4112709.552 | 0.00035827 |
| ML tree Modified (Bigyromonada+ochrophytes) | **0.0492** | -4112783.641 | 74.089 |

Except, the unconstrained ML tree, each tree was constrained under LG+C60+F+G4 using IQ-TREE with the dataset processed with approach 2. All the ML tree generated in this study (bigyromonada + oomycetes). “ML tree Modified” is a hypothetical tree constraint containing (bigyromonada + ochrophytes) with the rest of topology remaining the same with the unconstrained ML tree. The unconstrained tree is based on ML tree reconstructed under LG+C60+F+G4+PMSF as presented in Fig.S1. The p-AU values were calculated using the AU test with 10,000 RELL bootstrap replicates, implemented in IQ-TREE. The maximum log likelihoods (logL) of each constrained and their differences (ΔlogL) compared to the unstrained tree are listed. Constraints with P-values lower than 0.05 are rejected, indicating confidence interval below 95% (marked bold).
