## Supplemental Figure 3 for "Monophyly of Diverse Bigyromonadea and their Impact on Phylogenomic Relationships Within Stramenopiles"

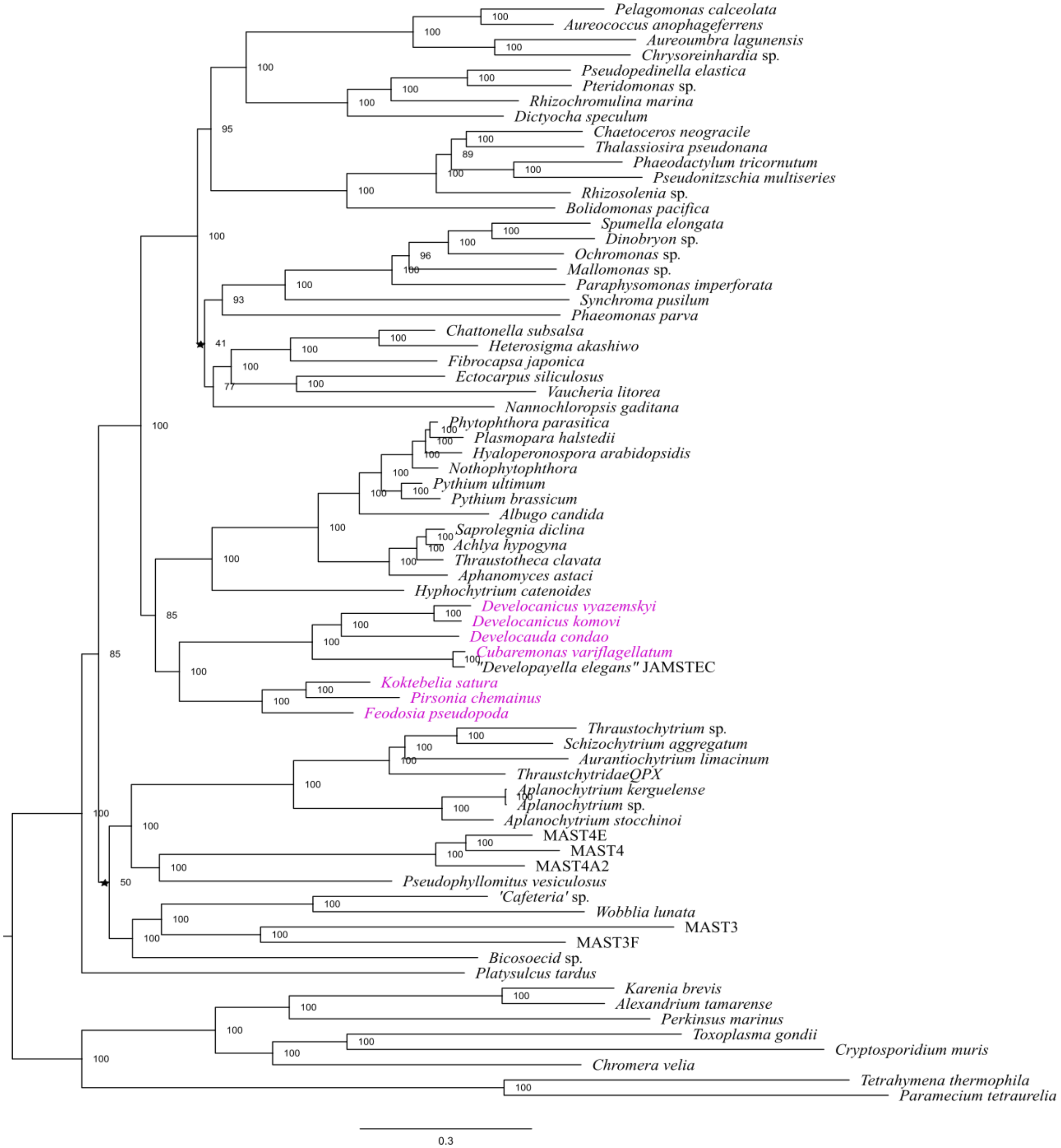


**Figure S3.** Multi-gene phylogenomic tree of stramenopiles with the seven new transcriptomes (pink) added to Gyrista with the fast-evolving species removed (*Cafeteria roenbergensis*, two species of *Blastocystis* sp., and *Cantina marsupialis*). The tree was reconstructed using the Maximum-likelihood (ML) analysis, under the site-heterogenous model (LG+C60+F+G4) implemented in IQ-Tree, comprising 75,798 aa of 247 genes from 72 taxa. Branch support was calculated using 1000 ultrafast bootstrap (UFB). Branches that have different topology from Fig. 1 are marked by a star symbol.
