## Supplemental Figure 4 for "Monophyly of Diverse Bigyromonadea and their Impact on Phylogenomic Relationships Within Stramenopiles"

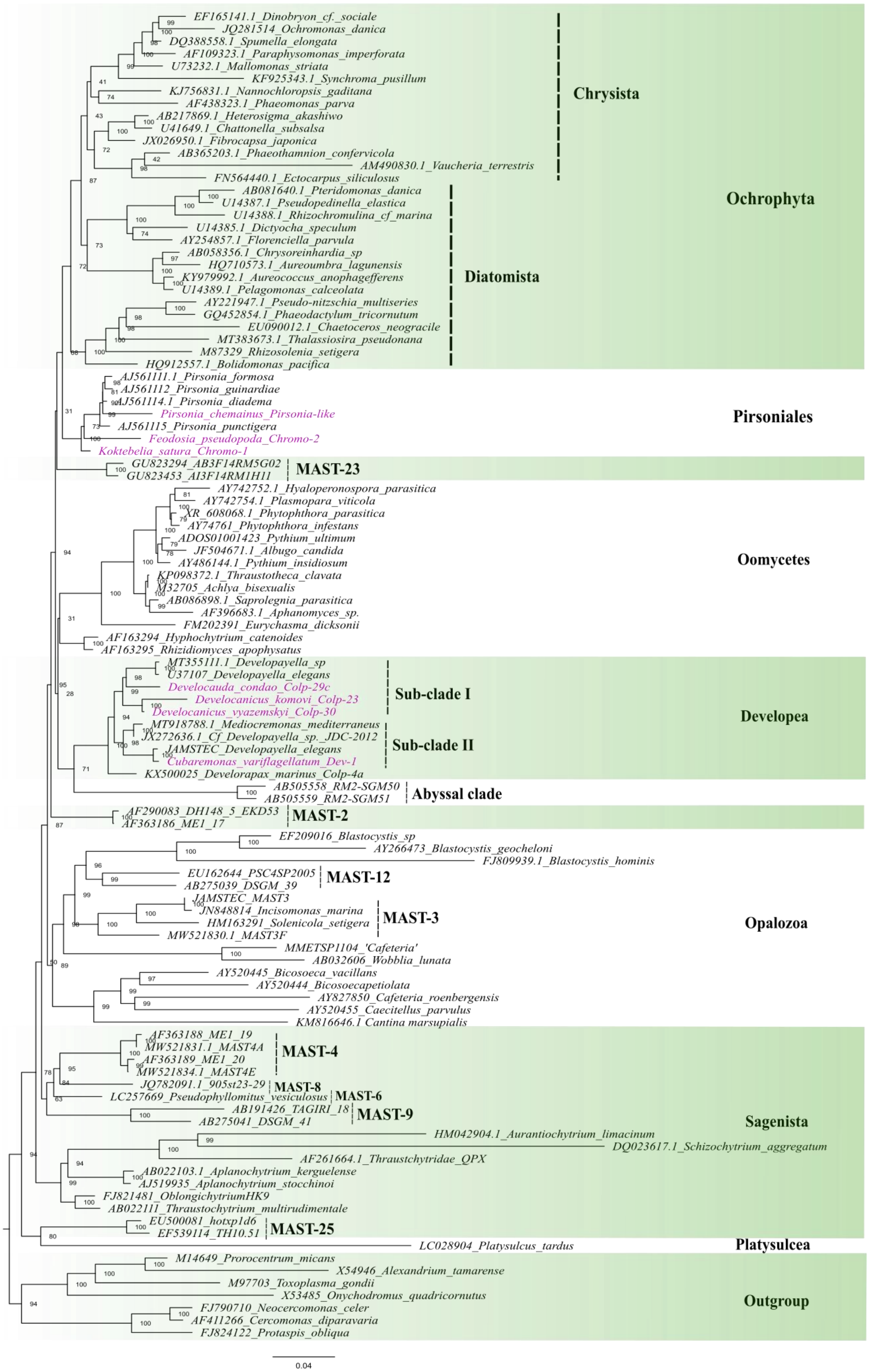
EF165141.1_Dinobryon_cf._sociale
JQ281514_Ochromonas_danica
DQ388558.1_Spumella_elongata
AF109323.1_Paraphysomonas_imperforata
U73232.1_Mallomonas_striata
KF925343.1_Synchroma_pusillum
KJ756831.1_Nannochloropsis_gaditana
AF438323.1_Phaeomonas_parva
AB217869.1_Heterosigma_akashiwo
U41649.1_Chattonella_subsalsa
JX026950.1_Fibrocapsa_japonica
AB365203.1_Phaeothamnion_confervicola
AM490830.1_Vaucheria_terrestris
FN564440.1_Ectocarpus_siliculosus
Chrysista
AB081640.1_Pteridomonas_danica
U14387.1_Pseudopedinella_elastica
U14388.1_Rhizochromulina_cf_marina
U14385.1_Dictyocha_speculum
AY254857.1_Florenciella_parvula
AB058356.1_Chrysoreinhardia_sp
HQ710573.1_Aureoumbra_lagunensis
KY979992.1_Aureococcus_anophagefferens
U14389.1_Pelagomonas_calceolata
AY221947.1_Pseudo-nitzschia_multiseries
GQ452854.1_Phaeodactylum_tricornutum
EU090012.1_Chaetoceros_neogracile
MT383673.1_Thalassiosira_pseudonana
M87329_Rhizosolenia_setigera
HQ912557.1_Bolidomonas_pacifica
Diatomista
Ochrophyta
AJ561111.1_Pirsonia_formosa
AJ561112_Pirsonia_guinardiae
AJ561114.1_Pirsonia_diadema
Pirsonia_chemainus_Pirsonia-like
AJ561115_Pirsonia_punctigera
Feodosia_pseudopoda_Chromo-2
Koktebelia_satura_Chromo-1
Pirsoniales
GU823294_AB3F14RM5G02
GU823453_AI3F14RM1H11
MAST-23
AY742752.1_Hyaloperonospora_parasitica
AY742754.1_Plasmopara_viticola
XR_608068.1_Phytophthora_parasitica
AY74761_Phytophthora_infestans
ADOS01001423_Pythium_ultimum
JF504671.1_Albugo_candida
AY486144.1_Pythium_insidiosum
KP098372.1_Thraustotheca_clavata
M32705_Achlya_bisexualis
AB086898.1_Saprolegnia_parasitica
AF396683.1_Aphanomyces_sp.
FM202391_Eurychasma_dicksonii
AF163294_Hyphochytrium_catenoides
AF163295_Rhizidiomyces_apophysatus
Oomycetes
MT355111.1_Developayella_sp
U37107_Developayella_elegans
Develocauda_condao_Colp-29c
Develocanicus_komovi_Colp-23
Develocanicus_vyazemskyi_Colp-30
Sub-clade I
MT918788.1_Mediocremonas_mediterraneus
JX272636.1_Cf_Developayella_sp._JDC-2012
JAMSTEC_Developayella_elegans
Cubaremonas_variflagellatum_Dev-1
Sub-clade II
KX500025_Developax_marinus_Colp-4a
Developea
AB505558_RM2-SGM50
AB505559_RM2-SGM51
Abyssal clade
AF290083_DH148_5_EKD53
AF363186_ME1_17
MAST-2
EF209016_Blastocystis_sp
AY266473_Blastocystis_geocheloni
FJ809939.1_Blastocystis_hominis
EU162644_PSC4SP2005
AB275039_DSGM_39
MAST-12
JAMSTEC_MAST3
JN848814_Incisomonas_marina
HM163291_Solenicola_setigera
MW521830.1_MAST3F
MAST-3
MMETSP1104_'Cafeteria'
AB032606_Wobblia_lunata
AY520445_Bicosoeca_vacillans
AY520444_Bicosoecapetiolata
AY827850_Cafeteria_roenbergensis
AY520455_Caecitellus_parvulus
KM816646.1 Cantina marsupialis
Opalozoa
AF363188_ME1_19
MW521831.1_MAST4A
AF363189_ME1_20
MW521834.1_MAST4E
MAST-4
JQ782091.1_905st23-29
MAST-8
LC257669_Pseudophyllomitus_vesiculosus
MAST-6
AB191426_TAGIRI_18
AB275041_DSGM_41
MAST-9
HM042904.1_Aurantiochytrium_limacinum
DQ023617.1_Schizochytrium_aggregatum
AF261664.1_Thraustchytridae_QPX
AB022103.1_Aplanochytrium_kerguelense
AJ519935_Aplanochytrium_stocchinoi
FJ821481_OblongichytriumHK9
AB022111_Thraustochytrium_multirudimentale
EU500081_hotxp1d6
EF539114_TH10.51
MAST-25
Sagenista
LC028904_Platysulcus_tardus
Platysulcea
M14649_Prorocentrum_micans
X54946_Alexandrium_tamarense
M97703_Toxoplasma_gondii
X53485_Onychodromus_quadricornutus
FJ790710_Neocercomonas_celer
AF411266_Cercomonas_diparavaria
FJ824122_Protaspis_obliqua
Outgroup
0.04

**Figure S4**. ML tree reconstructed under BIC: TIM2+R6 with 1000 UFB from a 18S rRNA gene alignment of 107 taxa (1665 sites) including environmental sequences. The seven new species described in this study are marked as pink: Pirsoniales forming a sisterhood with Ochrophytes and Developea forming a sister clade to ‘Abyssal Clade’, demonstrating potential expansion of the Bigyromonada clade with further taxon sampling.
